# Pan-flavivirus analysis reveals sfRNA-independent, 3’UTR-biased siRNA production from an Insect-Specific Flavivirus

**DOI:** 10.1101/2022.08.18.504478

**Authors:** Benoit Besson, Gijs J. Overheul, Michael T. Wolfinger, Sandra Junglen, Ronald P. van Rij

**Affiliations:** Department of Medical Microbiology, Radboud University Medical Center, Nijmegen, The Netherlands; Research Group Bioinformatics and Computational Biology, Faculty of Computer Science, University of Vienna, Vienna, Austria; Department of Theoretical Chemistry, University of Vienna, Vienna, Austria; RNA Forecast e.U.,Vienna, Austria; Institute of Virology, Charité-Universitätsmedizin Berlin, Corporate Member of Free University, Humboldt-University and Berlin Institute of Health, Berlin, Germany

**Author notes:** Correspondence to Ronald P. van Rij. Benoit Besson, Université de Strasbourg, RNA architecture and reactivity & Insect Models of Innate Immunity, Institut de Biologie Moléculaire et Cellulaire, CNRS, Strasbourg, France. Benoit Besson and Gijs J Overheul contributed equally to this work. Author order was determined alphabetically.

## Abstract

RNA interference (RNAi) plays an essential role in mosquito antiviral immunity, but it is not known whether viral siRNA profiles differ between mosquito-borne and mosquito-specific viruses. A pan-Orthoflavivirus analysis in *Aedes albopictus* cells revealed that viral siRNAs were evenly distributed across the viral genome of most representatives of the *Flavivirus* genus. In contrast, siRNA production was biased towards the 3’ untranslated region (UTR) of the genomes of classical insect-specific flaviviruses (cISF), which was most pronounced for Kamiti River virus (KRV), a virus with a unique, 1.2 kb long 3’ UTR. KRV-derived siRNAs were produced in high quantities and almost exclusively mapped to the 3’ UTR. We mapped the 5’ end of KRV subgenomic flavivirus RNAs (sfRNAs), products of the 5’-3’ exoribonuclease XRN1/Pacman stalling on secondary RNA structures in the 3’ UTR of the viral genome. We found that KRV produces high copy numbers of a long, 1017 nt sfRNA1 and a short, 421 nt sfRNA2, corresponding to two predicted XRN1-resistant elements. Expression of both sfRNA1 and sfRNA2 was reduced in *Pacman* deficient *Aedes albopictus* cells, however, this did not correlate with a shift in viral siRNA profiles. We suggest that cISFs and particularly KRV developed a unique mechanism to produce high amounts of siRNAs as a decoy for the antiviral RNAi response in an sfRNA-independent manner.

**IMPORTANCE:** The *Flavivirus* genus contains diverse mosquito viruses ranging from insect-specific viruses circulating exclusively in mosquito populations to mosquito-borne viruses that cause disease in humans and animals. Studying the mechanisms of virus replication and antiviral immunity in mosquitoes is important to understand arbovirus transmission and may inform the development of disease control strategies. In insects, RNA interference (RNAi) provides broad antiviral activity and constitutes a major immune response against viruses. Comparing diverse members of the *Flavivirus* genus, we found that all flaviviruses are targeted by RNAi. However, the insect-specific Kamiti River virus was unique in that small interfering RNAs are highly skewed towards its uniquely long 3’ untranslated region. These results suggest that mosquito-specific viruses have evolved unique mechanisms for genome replication and immune evasion.

## INTRODUCTION

The *Orthoflavivirus* genus constitutes diverse phylogenetic clades of viruses, found in vertebrates and arthropods including mosquitoes (1). Mosquito-borne arboviruses are transmitted horizontally between mosquitoes and vertebrates, whereas insect-specific flaviviruses (ISF) are thought to be primarily transmitted vertically and restricted to their arthropod hosts (2, 3). ISFs are further separated into two distinct phylogenetic clades: lineage I or classical ISFs (cISF), a clade that branches at the base of the *Orthoflavivirus* genus, and lineage II or dual-host affiliated ISFs (dISF) that forms a separate phylogenetic clade embedded in vector-borne clades (4–6). While the healthcare and economic burden of arboviruses is well established (7), ISFs have been proposed as modulators of arbovirus transmission and are being explored for biotechnological applications such as vaccine development (8–10).

Flaviviruses have an ∼11 kb long, positive-sense genomic RNA ((+)gRNA), which circularizes via long range RNA-RNA interactions between their 5’ and 3’ untranslated regions (UTR) for RNA translation and replication (11, 12). Asymmetric replication is mediated via an antigenomic negative-sense RNA intermediate ((-)gRNA), which serves as a template for replication of the (+)gRNA and is hypothesized to be annealed either to its template and/or to newly synthesized (+)gRNA, forming double-stranded RNA (dsRNA) (13, 14).

Flaviviruses take advantage of the ability of RNA to form regulatory, evolutionarily conserved elements to produce a highly structured subgenomic flavivirus RNA (sfRNA) (15–18). Formation of sfRNA is regulated by exoribonuclease-resistant RNA (xrRNA) structures in the 3’ UTR, which typically encompass three-way junctions (3WJ) or stem-loop (SL) elements that adopt a particular fold, mediated by a pseudoknot (19). The tight and complex structure of xrRNAs stalls the 5’-3’ exoribonuclease 1 (XRN1), also referred to as Pacman in mosquitoes, and terminates the degradation of viral RNA (20), resulting in the production of sfRNA. Flaviviruses may encode multiple xrRNA-like structures (21), each of which can induce the production of a distinct sfRNA species. While the longest sfRNA generated from the first xrRNA is generally the most abundant, sfRNA production from individual xrRNAs may vary between mammalian and mosquito hosts, suggesting viral adaptation to the host (20, 22–24).

It is well established that sfRNA is essential for flavivirus replication and dissemination (16, 23, 25, 26), for which several mechanisms have been suggested, in some cases with sfRNA serving as a decoy for the viral genome. For example, sfRNA was shown to inhibit the host RNA decay pathway (27), to control apoptosis (16, 28), to encode a microRNA (29), and to inhibit the mosquito Toll pathway (30). Moreover, sfRNA can be a substrate for small interfering RNA (siRNA) production by Dicer (31) and was proposed to inhibit the RNA interference (RNAi)-based antiviral immune response (27, 32–34), although this was recently disputed (28).

Mosquitoes have an RNAi-centered immune response, and deficiency in RNAi leads to increased sensitivity to virus infections (35–41). Viral dsRNA is cleaved by Dicer-2 into 21 nt viral siRNA duplexes (vsiRNAs), which are loaded into the Argonaute-2-containing RISC complex with the help of RNA-binding proteins Loqs and R2D2 (38, 42). Upon loading the duplexes, one of the RNA strands (passenger strand) is degraded and the remaining guide strand is used by Argonaute-2 to recognize and cleave complementary single-stranded viral RNA.

In addition to the siRNA pathway, the PIWI-interacting RNA (piRNA) pathway has been implicated in antiviral defense in mosquitoes (38, 43, 44). In this pathway, viral single-stranded RNA is processed into mature 25–30 nt viral piRNAs (vpiRNAs) associating with the PIWI proteins Piwi5 and Ago3, which amplify the piRNA response in a feedforward mechanism called the ping-pong amplification loop (45–47). While Piwi5 is required for vpiRNA biogenesis in *Aedes aegypti* (45, 46), only *Piwi4* depletion has thus far been shown to affect arbovirus replication (48, 49) and the importance of the piRNA response during acute viral infections remains to be clarified. Yet, endogenous viral elements (EVE) in the genome of *Aedes* mosquitoes give rise to piRNAs that can target cognate viral RNA and reduce viral RNA levels, especially in the ovaries (49–52), underlining the antiviral potential of the piRNA pathway.

While the RNAi response seems to be a widely active antiviral response, it is currently unclear whether mosquito-borne viruses and insect-specific viruses are differentially targeted. To address this question, we profiled small RNAs in *Aedes albopictus* mosquito cells infected with mosquito-borne viruses and ISFs and found that while siRNAs mapped across the entire length of the viral genome for all mosquito-borne flaviviruses tested, vsiRNAs were predominantly derived from the 3’ UTR of Kamiti River virus (KRV), a cISF originally identified in *Aedes mcintoshi* mosquitoes (53). A similar, but less pronounced trend was also observed for two other cISFs, Culex flavivirus (CxFV) and cell-fusing agent virus (CFAV). Having noted that KRV has a particularly long 3’ UTR, we set out to characterize viral subgenomic RNA species and mapped two main sfRNAs regulated by two predicted XRN1/Pacman-resistant xrRNA structures. Loss of *Pacman* resulted in a shift in sfRNA production, but no concomitant shift in siRNA profiles, suggesting that the 3’UTR biased siRNAs are produced in an sfRNA independent manner. We speculate that KRV and likely other cISFs developed a unique mechanism to evade antiviral RNAi.

## RESULTS

### RNAi response to flavivirus infection in *Aedes* mosquito cells

Given the importance of the siRNA response for antiviral immunity in mosquitoes, we analyzed viral small RNAs produced during flavivirus infection. Representatives of each major clade of mosquito-associated flaviviruses were selected to provide a pan-flavivirus overview of viral siRNA and piRNA profiles in *Ae. albopictus* U4.4 cells. Specifically, we selected the *Culex*-associated arboviruses Saint-Louis encephalitis virus (SLEV, isolate MSI-7) and West Nile virus (WNV), the *Aedes*-associated arboviruses dengue virus (DENV serotype 2) and Zika virus (ZIKV), the dISF Nounané virus (NOUV), the *Culex*-associated cISF CxFV, and the *Aedes*-associated cISFs CFAV and KRV (Fig. 1A). Further, the epidemic SLEV MSI-7 strain was compared to the ancestral SLEV Pal strain as representatives of cosmopolitan and enzootic viruses, respectively (54). All tested flaviviruses replicated to similar levels in U4.4 cells with approximately 10^8^ RNA copies/µg of total RNA at 72 hours post infection, except for CxFV and CFAV, which reached 3–5.10^6^ copies/µg of total RNA (Fig. S1).

**Figure 1.**
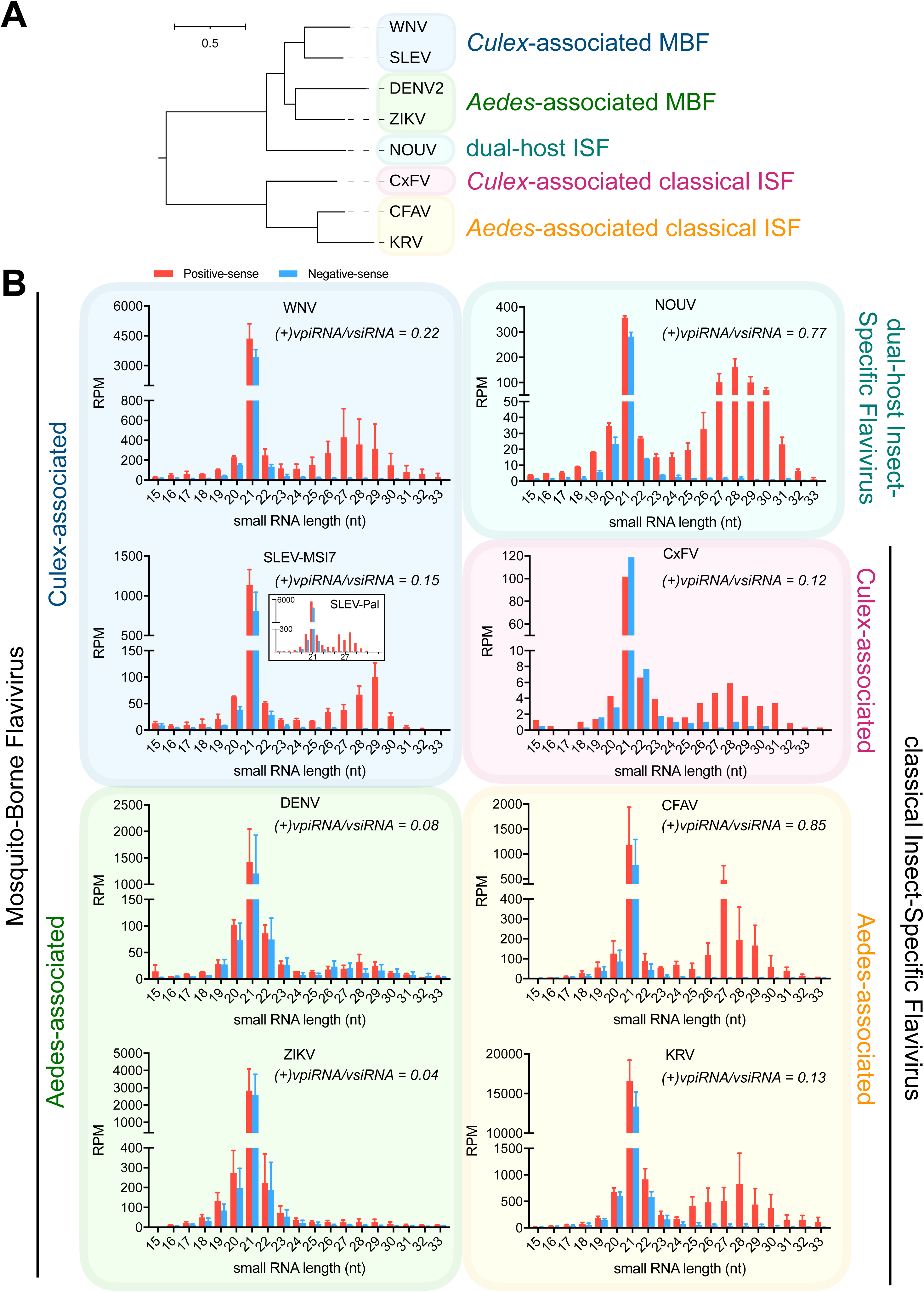
Comparison of flavivirus-derived small RNAs in U4.4 cells. **(A)** Maximum likelihood phylogenic tree based on whole genome sequences of the indicated viruses. Branch lengths are proportional to the number of substitutions per site. MBF, mosquito-borne flavivirus; ISF, insect-specific flavivirus. **(B)** Size profiles of flavivirus-derived small RNAs in read per million (RPM) from U4.4 cells infected for 72 h at an MOI of 0.1. Positive-sense RNAs are shown in red, negative-sense RNA in blue. Ratios of viral piRNAs over siRNAs are indicated for each virus. The results are the average of two experiments for all flaviviruses, except for CxFV (n = 1), CFAV (n = 3, of which 2 in WT U4.4 cells and 1 in CRISPR CTRL U4.4 cells), and KRV (n = 3). Error bars are the standard deviation between replicates.

As observed previously (3, 31, 45, 48, 50, 55, 56), size profiles of viral small RNAs were characterized by a prominent peak of 21 nt vsiRNAs from both positive- and negative-sense RNA for all tested flaviviruses (Fig. 1B), with a shoulder of predominantly positive-sense RNAs of 25–30 nt. Given similar viral RNA levels (Fig. S1A), differences in scales suggest that NOUV elicited an overall weaker siRNA response compared to WNV, SLEV-MSI-7/Pal, DENV and ZIKV. CxFV also elicited a weaker siRNA response, which may be due to the lower RNA levels (Fig. S1B). In contrast, KRV elicited the strongest siRNA response recorded, and CFAV induced a strong siRNA response despite its relatively low RNA levels in cells.

Although we have not formally demonstrated PIWI-protein association, the shoulder of 25-30 nt viral small RNAs (Fig. 1B) likely represent vpiRNAs. Indeed, the 1U-bias characteristic of PIWI-associated piRNAs was detectable for 25–30 nt sized RNAs of CFAV, KRV, DENV and SLEV (Fig. S2A). Moreover, a 10A-bias characteristic of piRNAs produced by ping-pong amplification was detectable on positive-sense, piRNA-sized RNAs of CFAV, KRV, DENV. Using the gRNA as reference, the flaviviruses differed from each other in the relative amounts of 25–30 nt viral small RNAs (Fig. S1C). Notably, viral piRNA over siRNA ratios were relatively low for the *Aedes*-associated arboviruses DENV and ZIKV (0.08 and 0.04, respectively), whereas these ratios were higher for NOUV and CFAV (0.77 and 0.85, respectively) (Fig. 1B).

### Asymmetric distribution of vsiRNAs across the KRV genome

The distribution of vsiRNAs across the viral genomes (Fig. 2) showed relative uniform mapping of vsiRNAs on both the (+)gRNA and (-)gRNA. A notable exception was KRV, for which most vsiRNAs mapped to the 3’ UTR region and presentation of the data on a logarithmic scale is required to show siRNAs mapping to other parts of the genome, albeit at extremely low levels (Fig. 2B). This pattern is reminiscent of 3’ UTR biased mapping observed for the other cISFs CxFV, CFAV and AEFV, although the skewed distribution is much more pronounced for KRV (Fig. 2A and C) (3, 50, 57). About 14% of the vsiRNAs of CxFV and CFAV and more than 95% of KRV vsiRNAs derived from their 3’ UTRs, in stark contrast to the other flaviviruses for which a median of ∼4% of vsiRNAs mapped to the 3’ UTR (Fig. 2C).

**Figure 2.**
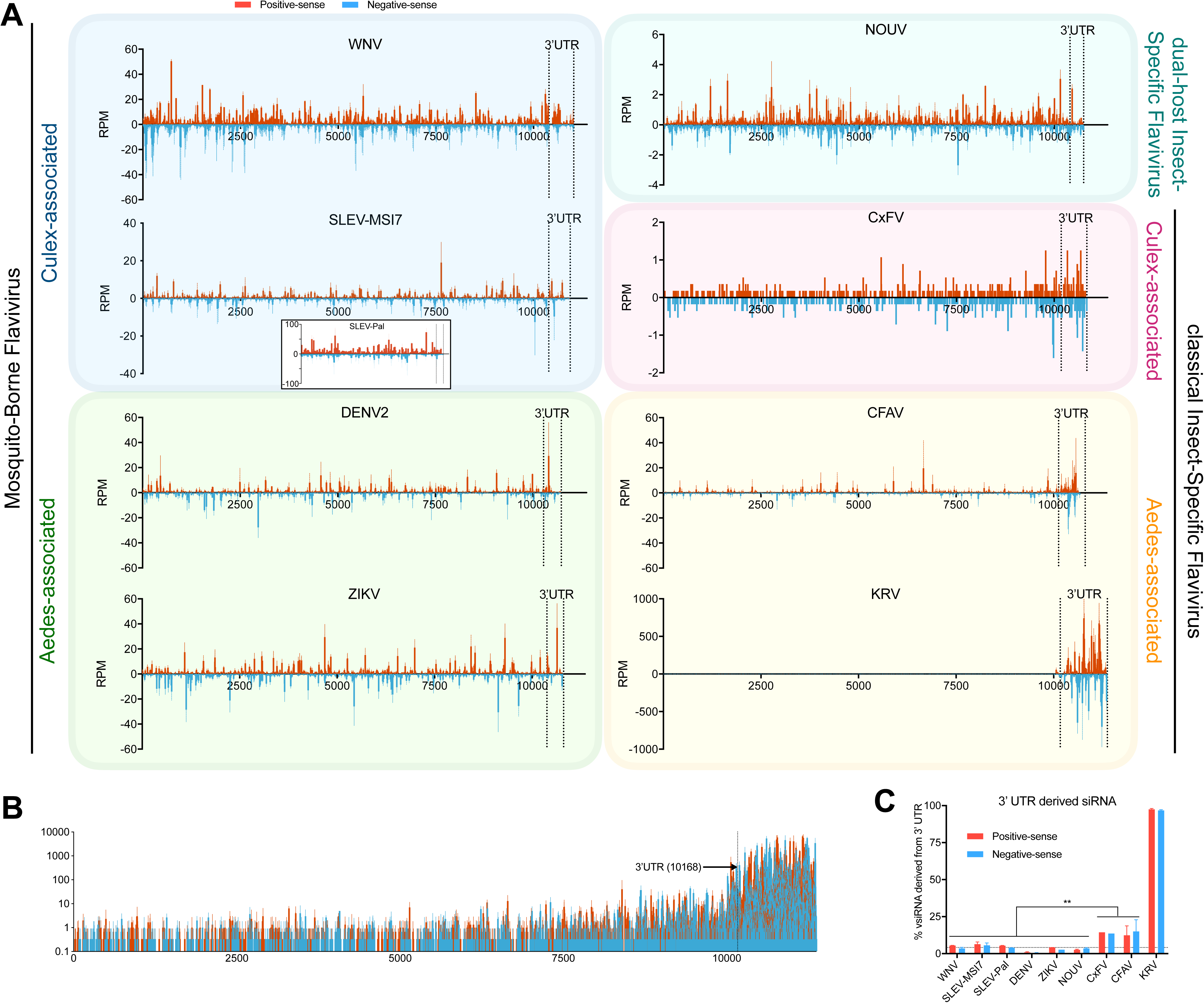
KRV vsiRNAs are strongly biased towards the 3’ UTR. **(A)** Distribution of flavivirus-derived vsiRNAs across the genome of each virus in reads per million (RPM) from U4.4 cells infected at a MOI of 0.1 for 72h. Start and end of the 3’ UTRs are indicated by dashed vertical lines. **(B)** Distribution of KRV vsiRNAs on a logarithmic scale with the position of the 3’ UTR indicated. **(C)** Percentage of vsiRNAs mapping to the 3’ UTR compared to the whole genome sequence for the indicated flaviviruses. The dashed horizontal line indicates the median of 3’ UTR derived vsiRNA from non-cISFs. (A–C) The results are the average of two experiments for all flaviviruses, except for CxFV (n = 1) and KRV (n = 3). Error bars are the standard deviation between replicates for each individual nucleotide. Positive-strand RNAs are shown in red, negative-strand RNAs in blue. ** *p* < 0.01; by two-way ANOVA and Fisher’s LSD test.

In contrast to siRNAs, vpiRNAs mapped to several discrete hotspots on the viral (+)gRNA (Fig. S2). For each virus analyzed, vpiRNAs mapped to different genome coordinates in a manner that was highly reproducible in replicate experiments, in agreement with previous observations (43, 48, 50, 58). It is worth noting that KRV derived piRNAs mapped at several hotspot across the gRNA and were not enriched at the 3’ UTR, indicating that each pathway processes a different substrate. Altogether, our data illustrate a general antiviral siRNA response to flaviviruses, no major differences between the ancestral and pandemic SLEV strains, and highlight the unique case of cISFs, especially KRV, for which the skewed distribution of vsiRNA towards the 3’ UTR suggests a unique siRNA response to the infection.

### KRV has a unique 3’ UTR

KRV has a 3’ UTR of 1208 nt, much longer than in any other member of the *Flavivirus* genus (median of 486 nt), but also longer that the 3’ UTRs of members of the cISF clade (median of 663 nt) (Fig. 3A). Structure predictions suggested that the KRV 3’ UTR is highly structured, comprising evolutionarily conserved elements, alongside secondary RNA structures that appear to be unique to KRV (Fig. 3B). Our model predicted the signature flavivirus regulatory SL at the 3’ end of the genome and corroborated the presence of two cISF xrRNA structures (xrRNA1 and xrRNA2) that are highly conserved between KRV, CFAV and Aedes Flavivirus (AEFV), which are closely related to *Anopheles*-associated cISFs (59, 60), and not conserved in the more distant Culex Flavivirus (CxFV) and Xishuangbanna Aedes Flavivirus (XFV). Moreover, structure predictions of the KRV 3’ UTR suggested the presence of simple and branched stem-loop elements, as well as several long hairpins, including the internal 3’ stem-loop (i3’SL), previously predicted using a comparative genomics approach (21).

**Figure 3.**
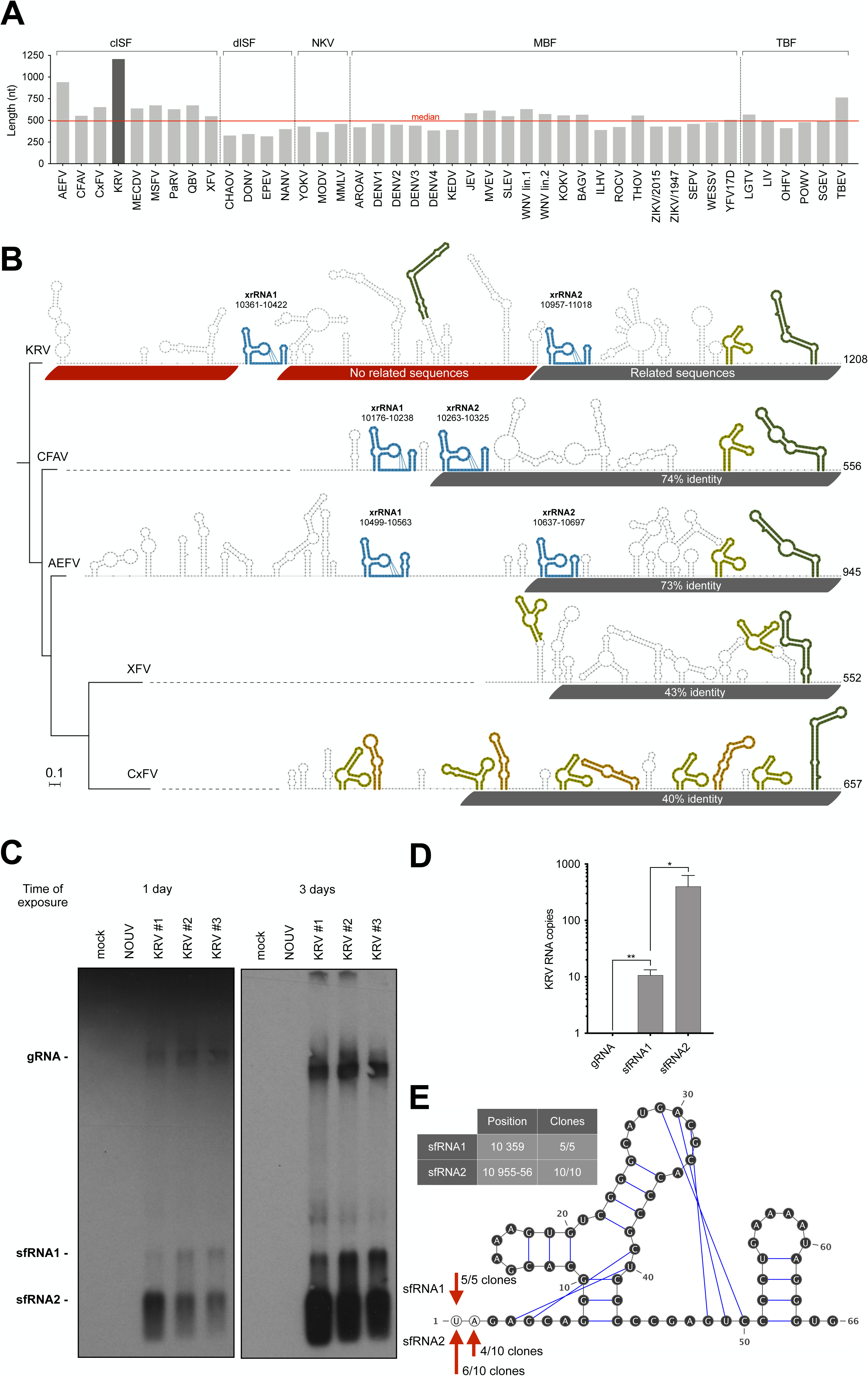
KRV has a long and unique 3’ UTR and produces high quantities of sfRNA. **(A)** Length of the 3’ UTR of all members of the *Flavivirus* genus with a RefSeq, a complete coding genome and a 3’ UTR of at least 200 nt. Viruses belong to the clades indicated: cISF, classical insect specific flaviviruses; dISF, dual-host affiliated insect specific flaviviruses; MBF, mosquito-borne flaviviruses; NKV, no known vector; TBF, tick-borne flaviviruses. For virus name and accession numbers, see Table S1. **(B)** Secondary structure prediction of the 3’UTR of four ISFs. Maximum likelihood phylogenic tree and alignment of 3’ UTR of listed viruses with conserved regions as described in (87). Branch lengths are proportional to the number of substitutions per site. Evolutionarily conserved RNA elements are highlighted in colour, with structurally homologous elements in the same colour. Elements without colour represent locally stable RNA structures from single-sequence RNA structure predictions. Exoribonuclease-resistant structures (xrRNA) in KRV, CFAV and AEFV are shown in blue, including reported pseudoknot interactions (17) with sequence regions downstream of the three-way junction structures. Repeat elements a (Ra) and b (Rb) (21) are depicted in olive and orange, respectively. 3’ stem-loop elements (3’SL) are shown in dark green. The internal 3’SL element of KRV is predicted to adopt a longer closing stem, which lacks evolutionary support in other viruses. The same applies for the extended closing stems of Ra elements in XFV. Percent nucleotide identities of each virus to KRV are indicated for the region between xrRNA2 and the 3’ SL. Lengths of the 3’ UTRs are indicated on the right. **(C)** Northern blot of positive-sense viral RNA in U4.4 cells infected with either NOUV or KRV at an MOI of 0.1 or mock infected for 72 h. Viral RNA was detected using a pool of oligonucleotide probes complementary to the 3’ UTR of KRV, between positions 10,361 and 11,375. All images were captured from the same northern blot. **(D)** Relative RT-qPCR quantification of KRV RNA in U4.4 cells infected for 72 h at an MOI of 0.1. Data are expressed relative to gRNA copy numbers and bars indicate means and standard deviation of four replicates. * *p* < 0.05; ** *p* < 0.01 by one-Way ANOVA and Fisher’s LSD test. **(E)** Position of 5’ ends of KRV sfRNA1 and sfRNA2 defined by 5’ to 3’ end ligation and sequencing, displayed on the consensus xrRNA structure predicted from SHAPE data for CFAV xrRNA1, which is 90% identical to KRV xrRNA1 and xrRNA2 (59).

Interestingly, while the 3’ terminal 419 nt long sequence of the KRV 3’ UTR downstream of xrRNA2 appears to be conserved with other cISFs (21), the 5’ terminal 789 nt sequence extending from the stop codon to xrRNA2 appears to be unique to KRV (Fig. 3B). This 5’ sequence of KRV 3’ UTR does not seem to share ancestry with AEFV, the cISF with the second longest 3’ UTR (Fig. 3A), nor with other flaviviruses, with the exception of xrRNA1 which is highly conserved both in structure and sequence, and was hypothesized to be the result of a self-duplication event (61, 62).

### KRV produces multiple subgenomic flavivirus RNA species

Given the long KRV 3’ UTR and the observation that flavivirus 3’ UTRs give rise to sfRNAs, we visualized the RNA species produced during KRV infection of *Ae. albopictus* U4.4 cells by northern blot (Fig. 3C left panel). Two sfRNA (sfRNA1 and sfRNA2) were detected, likely the product of XRN1/Pacman stalling on xrRNA structures. Both sfRNAs were displayed strong signals corresponding to the expected sizes (∼1000 and ∼400 nt), suggesting that KRV sfRNA1 and sfRNA2 greatly outnumber KRV (+)gRNA, as observed for other flaviviruses as well (16, 28, 31). When quantifying the different KRV RNA species by RT-qPCR (Fig. 3D), we found that for each molecule of gRNA, there was a 10-fold increase of sfRNA1 and 400-fold increase of sfRNA2 relative to gRNA, confirming the presence and high abundance of the two sfRNAs during KRV infection. Finally, we characterized the 5’ start site of the subgenomic RNA species using a 5’-3’ ligation assay. This analysis confirmed that sfRNA1 and sfRNA2 started immediately upstream of xrRNA1 and xrRNA2, resulting in products of 1017 nt and 421-422 nt, respectively (Fig. 3E).

With the new characterization of the KRV sfRNAs, we further analyzed our small RNA sequencing data (Fig. 1–2) and investigated the vsiRNA distribution on the genomic region corresponding to the sfRNA downstream from the xrRNA structures, either expressed as a percentage of the genome-mapping vsiRNAs (Fig. 4A) or as a density of vsiRNA reads per nt (Fig. 4B). The sfRNA region of CFAV, which covers 92% of its 3’UTR (Fig. 3B), was clearly associated with a higher density of vsiRNAs. In contrast, only a negligible amount of KRV vsiRNA derived from its gRNA-specific regions (<7%), whereas 92% of KRV vsiRNA mapped to the sfRNA region in the 3’ UTR. Thus, the 3’ bias characteristic of cISF and especially KRV-derived vsiRNAs correlated highly with the sfRNA region of the 3’UTR.

**Figure 4.**
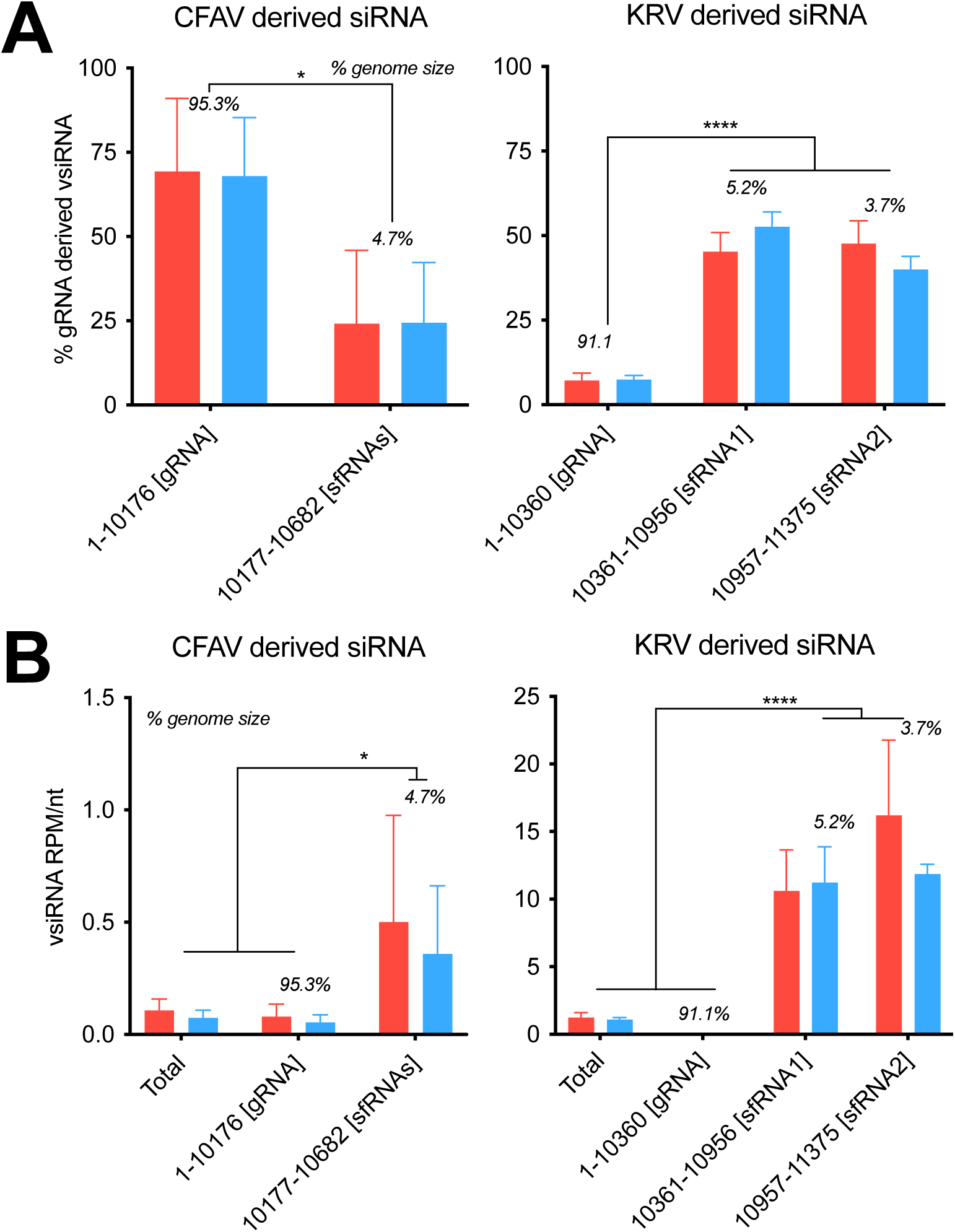
cISF-derived siRNA correlate with the sfRNA region of their genome. **(A)** Percentage of vsiRNAs compared to the whole genome sequence and **(B)** average density of vsiRNAs per nucleotide in the indicated regions of the genome of CFAV (left) or KRV (right). The size of each region is indicated in italic as a percentage of the genome size. (A– B) The results are the average of two experiments for all flaviviruses, except for CxFV (n = 1) and KRV (n = 3). Error bars are the standard deviation between replicates for each individual nucleotide. Positive-strand RNAs are shown in red, negative-strand RNAs in blue. ** *p* < 0.01; **** *p* < 0.0001 by two-way ANOVA and Fisher’s LSD test.

### Biogenesis of KRV sfRNA is Pacman-dependent

To determine whether sfRNA biogenesis is *Pacman*-dependent, we used CRISPR/Cas9 gene editing to create *Pacman* knockout (KO) U4.4 cell lines. Several putative *Pacman* loci are annotated in the *Ae. albopictus* genome, of which AALFPA_065179, AALFPA_057530 and AALFPA_079140 contain the conserved 5’-3’ exoribonuclease domain (> 98% identity across loci), whereas AALFPA_052256 only contains the SH3-like domain and is unlikely to encode a functional Pacman nuclease (Fig. S3A). Guide RNAs were designed to introduce frameshift mutations leading to premature stop codons in the 5’-3’ exoribonuclease domain. Two *Pacman* KO U4.4 cell clones were obtained (g3#3 and g2#13), which were compared to a CRISPR control line (CTRL) that was subjected in parallel to the same treatment without functional guide RNA, and to the wildtype (WT) parental U4.4 cell line. *Pacman* mRNAs containing the 5’-3’ exoribonuclease domain were unstable in both *Pacman* KO U4.4 cell clones (Fig. S3B), likely due to nonsense mediated decay induced by the presence of premature stop codons. KRV replicated to similar levels in *Pacman* KO cells as in WT and CTRL cells (Fig. S3C).

Using northern blotting, we observed lower levels of sfRNA1 and sfRNA2 in KRV infected *Pacman* KO cells, confirming that their biogenesis is *Pacman*-dependent (Fig. 5A). Interestingly, two different ∼800 nt and ∼500 nt subgenomic RNAs were identified in *Pacman* KO cells, which we named sfRNA1’ and sfRNA2’, likely the products of redundant 5’-3’ exoribonucleases stalling on structures downstream of xrRNA1 (63, 64). This is reminiscent of the appearance of other RNA species without loss of sfRNA upon knockdown of XRN1 in human cells (65). Processing of viral RNA by other 5’-3’ exoribonucleases would also explain why viral gRNA was not stabilized in *Pacman* KO cells (Fig. 5A, Fig. S3C)

**Figure 5.**
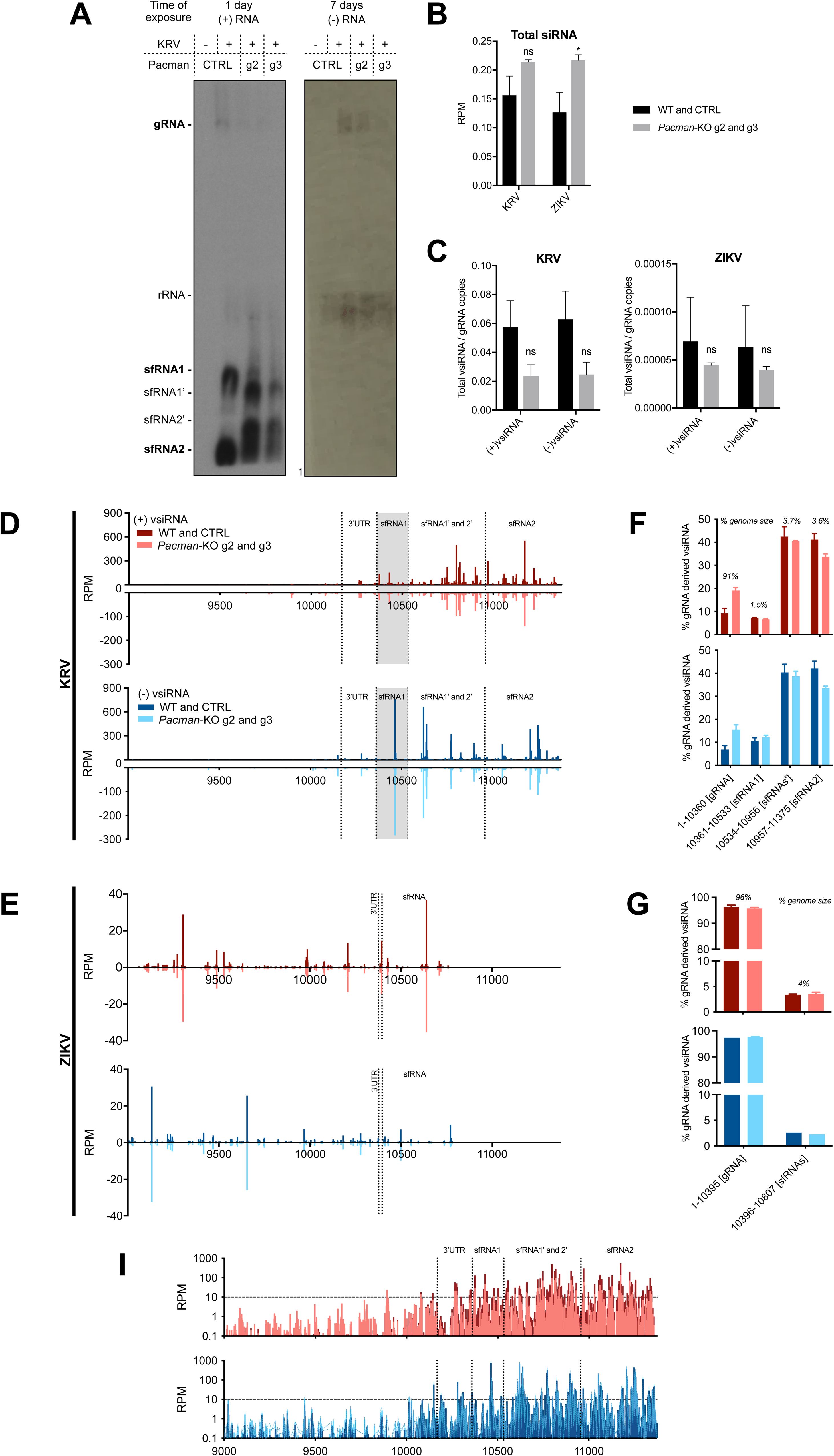
Pacman knockout does not affect vsiRNA profiles. (**A**) Northern blot of positive-sense ((+) RNA) or negative-sense ((-) RNA) viral RNA in control (CTRL) cells or *Pacman* knockout cells (clones g3#3 and g2#13), mock infected (-) or infected with KRV (+) at an MOI of 0.1 for 72 h. Viral RNA was detected using a pool of oligonucleotide probes complementary to the 3’ UTR of KRV, between positions 10,361 and 11,375. The location of ribosomal RNA (rRNA) based on ethidium bromide staining is indicated. **(B–C)** Quantification of total siRNAs (B) and sense (+) and antisense (-) vsiRNAs (C) in wild-type (WT), CRISPR control (CTRL), or *Pacman* knockout (KO) U4.4 cell lines infected with ZIKV at an MOI of 0.1 or KRV at an MOI of 10 for 72 h. Viral siRNA levels were normalized to viral gRNA levels. Errors bars represent the standard deviation from two independent cell lines. ns, non-significant; *, *p* < 0.05 by two-way ANOVA and Fisher’s LSD test. **(D-E)** Distribution of (+) and (-) vsiRNAs across the 3’ end of the genomes of KRV (**D**) or ZIKV (**E**) (from nt 9000 onwards) in CTRL and *Pacman* KO cells. Top panels show (+) vsiRNAs and lower panels show (-) vsiRNAs. The results are the average from two independent cell lines. **(F-G)** Percentage of vsiRNAs derived from the indicated regions compared to the entire gRNA. The size of each region is indicated in *italic* as a percentage of the viral genome size. Errors bars represent the standard deviation from two independent cell lines. (**I**) Distribution of KRV vsiRNAs on a logarithmic scale plotted on the 3’terminal region of the genome. (**D-E, I**) Boundaries of 3’ UTRs and sfRNAs are indicated by dashed vertical lines. (**D-G**) Blue, (+) vsiRNA; red, (-) vsiRNA; darker, WT and CTRL cells; lighter, *Pacman*-KO cells.

The exact 5’ start sites of sfRNA1’ and sfRNA2’ were determined by 5’-3’end ligation to be at nt 10,533 (9/9 clones) and 10,830 (4/5 clones) of the KRV genome, respectively. These sites did not correspond to notable predicted structures or RNA motifs (Fig. 3B, data not shown). Overall, the production of KRV sfRNA is *Pacman*-dependent and its absence leads to the production of alternative sfRNAs shorter than sfRNA1.

### Viral small RNA production in *Pacman* knockout cells

In absence of an infectious clone required to generate KRV without sfRNA and study its role in siRNA production, we took advantage of the shift in sfRNA production in *Pacman* KO cells as a substitute to investigate putative sfRNA-derived siRNAs. We explored the involvement of sfRNAs in the 3’ bias of KRV vsiRNAs by comparing vsiRNA profiles in U4.4 CTRL cells with *Pacman* KO cells, in which two new KRV subgenomic RNAs were produced (sfRNA1’ and 2’; Fig. 5A).

Interestingly, total siRNA levels were higher in *Pacman* KO cells than in control cells (Fig. 5B), perhaps due to the higher processing of mRNA by the siRNA pathway when the RNA decay pathway was impaired. In contrast, vsiRNA levels decreased slightly for KRV and ZIKV in the absence of Pacman (Fig. 5C, S3C). Surprisingly, in the segment differentiating KRV sfRNA1 from the *Pacman* KO associated sfRNA1’ and 2’ (nt 10361-10533), no difference in vsiRNA distribution was observed (Fig. 5D, 5F). Similarly, no major differences were observed for vsiRNA profiles of ZIKV between *Pacman* KO and CTRL cells (Fig. 5E, 5G). Further, presentation of the KRV vsiRNA on a logarithmic scale indicates that the 3’bias in starting upstream of the 3’UTR, around positions 9890-10010, in a *Pacman*-independent manner (Fig. 1B, 5I). These results demonstrate that while KRV 3’ biased vsiRNAs correlate strongly with the 3’UTR and by extension its sfRNA region, they are not produced from sfRNA species.

## DISCUSSION

Within the *Flavivirus* genus, cISFs constitute a unique clade of viruses that evolved independently, only infecting invertebrate hosts in which they are not associated with known disease (66). As such, cISFs represent a prime resource to better understand viral infection and antiviral immunity in mosquitoes. In a pan-flavivirus small RNA analysis, we found that vsiRNAs generally mapped across the viral genome for most mosquito-specific and mosquito-borne viruses, while there was a strong vsiRNA bias toward the 3’ UTR of KRV, the production of which was independent of its two, highly abundant sfRNAs.

### Unique RNAi response toward classical insect specific flaviviruses is sfRNA-independent

RNAi is a cornerstone of mosquito immunity comparable to the importance of the interferon response in mammalian systems, as its deficiency leads to increased sensitivity to viral infections (36, 67–69). Our analysis strengthens previous observations in *Anopheles* (70), *Culex* (55, 71) and *Aedes* (45, 67) that mosquito RNAi raises a broad and uniform antiviral response against all assessed mosquito-borne flaviviruses. Yet, cISFs seem to have evolved to produce a unique RNAi response with vsiRNAs biased towards the 3’ UTR of the viral genome, which was particularly strong for KRV but also detectable for CxFV, CFAV, and previously for AEFV (3). Interestingly, we did not observe a 3’ vsiRNA bias for the dISF NOUV, indicating that the biased vsiRNA production is not required for a mosquito-restricted transmission cycle.

The homogeneous distribution across the genome and the absence of a strand bias of vsiRNAs is consistent with processing of flaviviruses dsRNA consisting of (+) and (-) gRNA hybrids (36). The 3’ vsiRNA bias of KRV and other cISFs suggest a correlation with viral RNA species produced specifically by cISFs, which remain to be elucidated. A 3’ vsiRNA bias has previously been suggested to be related to sfRNA and RNA structure of the region (3), but our data do not support such hypothesis. First, KRV vsiRNAs derive equally from sfRNA1 or sfRNA2 regions, while more vsiRNAs would be expected toward the 3’ end due to the high abundance of both sfRNA1 and sfRNA2 (23, 28, 31). Second, a shift in sfRNA production is observed in *Pacman*-knockout cells, but this did not affect vsiRNA patterns. Third, (+) and (-) sense vsiRNAs are present at roughly equimolar levels, whereas sfRNA is a (+) sense RNA, which would result in a strong (+) strand bias of siRNAs should sfRNA be the Dicer-2 substrate. Fourth, the vsiRNA 3’ bias starts upstream of the 3’ UTR and the sfRNAs. Thus, the mechanism underlying the 3’ bias of vsiRNAs remains to be understood and may be multi-factorial. It was proposed that sfRNAs could bind the 5’ end of (-) gRNA as a competitive regulator of RNA replication (72–74), which would generate a double-stranded Dicer substrate, but this is not be consistent with our observations and arguments as outlined above, and would not account for a significant fraction of 3’UTR derived vsiRNAs. Therefore, previous suggestions that cISF 3’ biased siRNAs can be attributed to the sfRNA (3, 50, 57) are likely due to the sfRNA region covering most of the 3’UTR and a more moderate 3’ UTR biased siRNA profile that does not allow enough resolution. Instead, our data suggest the presence of an as yet unknown subgenomic dsRNA species of approximately 1500 nt that is colinear with the 3’ end of the viral genome. Effective processing of such a dsRNA molecule by the RNAi machinery may explain that it has escaped detection in our northern blot analyses.

## Conclusion

As part of the constant arms race between viruses and their hosts (75), cISFs and especially KRV seem to have evolved unique ways to maintain themselves in mosquito populations. KRV’s strikingly long 3’ UTR representing 10% of its total gRNA, combined with the expression of two highly abundant sfRNAs, as well as the strong 3’ bias of vsiRNAs makes KRV an intriguing model to study the biology of cISFs and the mechanisms of mosquito antiviral immunity.

## ACKNOWLEDGEMENTS

We thank Pascal Miesen for preliminary analysis of small RNA sequencing data. We thank Andrea Gamarnik (Fundación Instituto Leloir, Buenos Aires, Argentina) for advice on the 5’-3’ ligation method. We thank Beate Kümmerer (University of Bonn) for providing DENV2 strain 16681 and M. Niedrig (Robert-Koch-Institute, Berlin) for providing WNV strain NY-99. CFAV (isolate Rio Piedras), CxFV (isolate Uganda08), KRV (isolate SR-75) and ZIKV (isolate H/PF/2013) were provided by Xavier de Lamballerie (Aix Marseille Université) through the European Virus Archive (EVAg), funded by the European Union’s Horizon 2020 programme. This study was financially supported by the Deutsche Forschungsgemeinschaft (DFG) within Priority Programme SPP1596 [grant number JU 2857/1-1 and JU2857/1-2 to SJ and grant number VA 827/1-1 and RI 2381/1-2 to RPvR] and by a European Research Council Consolidator Grant under the European Union’s Seventh Framework Programme [grant number ERC CoG 615680 to RPvR]. Small RNA sequencing was performed by the GenomEast platform, a member of the “France Génomique” consortium (ANR-10-INBS-0009).

## MATERIAL AND METHODS

### Cells and viruses

*Aedes albopictus* C6/36 cells (ECACC General Cell Collection, #89051705) and U4.4 cells (kindly provided by G.P. Pijlman, Wageningen University, the Netherlands) were cultured at 28°C in Leibovitz L15 medium (Gibco) supplemented with 10% heat inactivated fetal calf serum (Sigma), 2% tryptose phosphate broth solution (Sigma), 1x MEM non-essential amino acids (Gibco), and 50 U/ml penicillin and 50 μg/ml streptomycin (Gibco).

CFAV (isolate Rio Piedras), CxFV (isolate Uganda08), KRV (isolate SR-75) and ZIKV (isolate H/PF/2013) were obtained from the European Virus Archive. DENV2 (strain 16681) was provided by Beate Kümmerer, University of Bonn. SLEV (strain MSI-7) was obtained from the National Collection of Pathogenic Viruses, Porton Down, Salisbury, United Kingdom and WNV (strain NY-99) was kindly provided by M. Niedrig, Robert-Koch-Institute, Berlin, Germany. SLEV (Palenque) and NOUV (isolate B3) were previously characterized (54, 76). Reference sequences are listed in Table S1.

### Plasmids

Plasmids used as standard to normalize the qPCR data were obtained by blunt end ligation of the flavivirus qPCR products into pGEM-3Z (Promega), using SmaI (NEB) and T4 DNA ligase (Roche). For KRV RNA species quantification, plasmid pUC57-KRV-9000-11375 containing part of NS5 gene and the 3’ UTR of KRV (nt 9000-11375) was synthesized by GenScript.

The pAc-Cas9-AalbU6.2 plasmid was generated by replacing the *D. melanogaster U6* promoter in the pAc-sgRNA-Cas9 (kindly provided by Ji-Long Liu, Addgene plasmid #49330) (77) with the *Ae. albopictus* promoter from AALFPA_045636 and by introducing XbaI restriction sites using In-Fusion cloning (Takara) on four fragments amplified using the primers in Table S2, according to the manufacturer’s instructions. sgRNA sequences targeting *Pacman* were cloned into a SapI restriction site directly downstream of the U6 promoter by annealing and phosphorylating complementary oligonucleotides (Table S2), followed by ligation using T4 DNA ligase (Roche), replacing the 5’-GGAAGAGCGAGCTCTTCC-3’ sequence that was used as negative control for CRISPR/Cas9 knockouts.

### CRISPR/Cas9

U4.4 cells were transfected with the pAc-Cas9-AalbU6.2 plasmid, which expresses 3xFLAG-tagged Cas9 with N- and C-terminal SV40 nuclear localization signals, followed by a viral 2A ribosome self-cleavage site and the puromycin N-acetyltransferase coding sequence, driven by the *D. melanogaster Actin5c* promoter. In parallel to the plasmid encoding *Pacman* sgRNA, cells were transfected with the parental pAc-Cas9-AalbU6.2 plasmid to generate the CRISPR control (CTRL) cells. U4.4 cells were seeded in a 24-well plate and transfected the next day with 500 ng of plasmid, using X-tremeGENE HP transfection reagent (Roche) according to the manufacturer’s instructions. At 2 days after transfection, puromycin (InvivoGen) was added to the culture medium at a concentration of 20 µg/ml and 4 days later, the cells were transferred to a new plate at a 1:2 dilution, in medium containing 20 µg/ml puromycin. The other half of the cells was used for genomic DNA isolation (Zymo Research #D3024) and PCR (Promega #M7806) using primers flanking the sequence targeted by the sgRNA (Table S2) to assess editing efficiency. Multiple sgRNA constructs were initially constructed and the constructs with the highest editing efficiency were selected, assessed by size heterogeneity of the PCR products on ethidium bromide-stained agarose gel. Cells transfected with these sgRNA constructs (g2 and g3) were seeded in 96-well plates at a density of a single cell per well in supplemented L15 medium in the absence of puromycin. After 3 weeks, gDNA was isolated from the single-cell clones, followed by PCR and Sanger sequencing of the targeted *Pacman* locus. Based on the sequencing results U4.4 clones g2#13 and g3#3, both containing only out-of-frame deletions in the *Pacman* coding region, were selected for further analyses.

### Small RNA library preparation and analysis

U4.4 cells were seeded at a density of 2×10^6^ cells per well in 6-well plates and infected the following day with the designated flaviviruses at an MOI of 0.1. Cells were harvested at 72 hours post infection (hpi) in RNA-Solv reagent (Omega Biotek R630-02) for total RNA isolation. 1 µg of RNA was used as input for library preparation using the NEBNext Multiplex Small RNA Library Prep Kit for Illumina (NEB E7560S), according to the manufacturer’s recommendations. The libraries were size-selected on 6% polyacrylamide/1x TBE gels, quantified using the Agilent 2100 Bioanalyzer System, and pooled for sequencing on an Illumina HiSeq4000 machine by the GenomEast Platform (Strasbourg, France). Viral small RNA sequences were mapped to the designated genome (Table S1) using Bowtie (Galaxy Tool Version 1.1.2) (78) allowing 1 mismatch. The genome distribution of the viral small RNAs was obtained by plotting the 5’ ends of mapping reads on the viral genome sequence. Nucleotide biases were plotted using the WebLogo 3 program (Galaxy Tool Version 0.4). All reads were normalized by library size as reads per million. The small RNA sequencing datasets have been deposited at the Sequence Read Archive under accession number PRJNA830662.

### Reverse transcription and quantitative PCR

For RT-qPCR, 1 µg of total RNA was reverse transcribed in a 20 µl reaction using Taqman RT Reagents (ThermoFisher Scientific) with hexamers at 48°C for quantification of viral RNA copies or SuperScript IV Reverse Transcriptase (ThermoFisher Scientific) with primer 5’-AGCGCATTTATGGTATAGAAAAGA-3’ at 60°C for quantification of specific KRV RNA species. Quantitative PCR was performed using the GoTaq qPCR SYBR mastermix (Promega) on a LightCycler 480 instrument (Roche). A standard curve of plasmids containing the corresponding viral sequence was used to convert Ct values to relative viral RNA copy numbers. *Pacman* mRNA levels were normalized to house-keeping gene *ribosomal protein L5*. For qPCR primer sequences, see Table S2.

### Northern Blot

5 µg of total RNA was separated on a 1X MOPS, 5% formaldehyde, 0.8% agarose gel for 5 h, transferred on a Hybond NX nylon membrane (Amersham) and cross-linked in the Gene linker UV chamber (Bio-Rad). Viral RNAs were detected with DNA oligonucleotides (Table S2) end-labelled with [^32^P] γ-adenosine-triphosphate (Perkin Elmer) using T4 polynucleotide kinase (Roche). Hybridization to the oligonucleotide probes was performed overnight at 42°C in Ultra-hyb Oligo hybridization buffer (Ambion). Membranes were then washed three times at 42°C with decreasing concentrations of SDS (0.2 to 0.1%) and exposed to X-ray films (Carestream).

### 5’ to 3’ end ligation

The 5’ ends of KRV RNA species were determined by 5’ to 3’ end ligation using a method adapted from (11, 79). C6/36 cells were seeded at a density of 4×10^6^ cells per T75 flask and infected the following day with KRV at an MOI of 10. Cells were harvested at 48 hpi and RNA was isolated using RNA-Solv reagent and 10 µg of total RNA was treated with T4 RNA ligase (Epicenter). Ligated RNAs were reverse transcribed with Taqman Multiscribe (Applied Biosystem) using random hexamers. The 5’-3’ junction region was amplified by PCR using a forward primer at the end of the 3’ UTR and a reverse primer in the 5’ part of KRV RNA species (Table S2), cloned into pGEM-3Z (Promega) using In-Fusion technology (Takara) and Sanger sequenced by the in-house sequencing facility.

### RNA structure prediction and bioinformatics

RNA secondary structure were predicted in the 3’UTR regions of KRV, CFAV, CxFV, and XFV using the ViennaRNA Package v.2.5.1 (80). Evolutionarily conserved elements were identified with the help of the viRNA GitHub repository (https://github.com/mtw/viRNA), and used as constraints for RNA structure prediction. Locally stable RNA structures were predicted with RNALfold from the ViennaRNA Package, allowing for a maximal base pair span of 100 nt.

Multiple sequence alignments of whole genome and 3’ UTR sequences were generated with MAFFT (81), curated with BMGE (82) and a maximum likelihood phylogenetic tree was built with PhyML (83) using NGPhylogeny.fr (84) using default settings. Phylogenetic trees were visualized on iTOL (85). Percentage identities were determined with Mview (86). Viral reference sequences are listed in Table S3.

### Statistical analysis

Graphical representation and statistical analyses were performed using GraphPad Prism v7 software. Differences were tested for statistical significance using one- or two-way ANOVA and Fisher’s LSD test.

---

**Supplementary figure 1. Relative quantification of gRNA, vsiRNA and vpiRNA.**

**(A)** Quantification of viral gRNA copies, **(B)** total vsiRNA over gRNA copy ratios, and **(C)** (+) vpiRNA over gRNA copy ratios in the samples analyzed in Figures 1–2 and S2. U4.4 cells were infected with the indicated virus at a MOI of 0.1 for 72h. Error bars are the standard deviation of at least 2 independent experiments. ns, non-significant by one-way ANOVA and Fisher’s LSD test.

**Supplementary figure 2. Viral piRNA profiles of flavivirus infected U4.4 cells** Distribution of flavivirus derived 25–30 nt vpiRNAs on the genome of each virus in reads per million (RPM) from U4.4 cells infected for 72h at a MOI of 0.1. In boxes, sequence logos at position 1 and 10 of 25–30 nt vpiRNAs. The results are the average of two experiments for all flaviviruses, except for CxFV (n=1) and KRV (n=3). Error bars are the standard deviation between replicates. Positive-strand RNAs are shown in red, negative-strand RNA in blue.

**Supplementary figure 3. Characterization of flavivirus infection of *Pacman*-KO U4.4 cells**

**(A)** *Pacman* KO U4.4 cell lines were generated by CRISPR/Cas9 technology, amplified from single clones, and the edited sites in exon 4 were Sanger sequenced. Sequencing identified three small deletions that all induced out-of-frame mutations in the exoribonuclease domain in both *Pacman* KO clones. Gene structure, conserved domains, and primer sets used in (B) are indicated. **(B)** Relative quantification by RT-qPCR of *Pacman* mRNA in CTRL and *Pacman* KO cells infected with KRV at a MOI of 1 for 96h. *Pacman* mRNA levels were normalized to house-keeping gene *ribosomal protein L5* and expressed relative to *Pacman* mRNA levels in WT cells. **(C)** Quantification of ZIKV or KRV RNA in cells and culture supernatants of the indicated U4.4 cells at 72 h post infection at an MOI of 0.1 or 10.

**Supplementary table 1. List of viruses used**

**Supplementary table 2. List of oligonucleotides for northern blots, qPCR and cloning**

**Supplementary table 3. List of genome references used for 3’UTR analysis**

## REFERENCES

1. Postler TS, Beer M, Blitvich BJ, Bukh J, de Lamballerie X, Drexler JF, Imrie A, Kapoor A, Karganova GG, Lemey P, Lohmann V, Simmonds P, Smith DB, Stapleton JT, Kuhn JH. 2023. Renaming of the genus Flavivirus to Orthoflavivirus and extension of binomial species names within the family Flaviviridae. Arch Virol 168:224.

2. Saiyasombat R, Bolling BG, Brault AC, Bartholomay LC, Blitvich BJ. 2011. Evidence of Efficient Transovarial Transmission of Culex Flavivirus by Culex pipiens (Diptera: Culicidae). J Med Entomol 48:1031–1038.

3. Frangeul L, Blanc H, Saleh M-C, Suzuki Y. 2020. Differential Small RNA Responses against Co-Infecting Insect-Specific Viruses in Aedes albopictus Mosquitoes. Viruses 12:468.

4. Blitvich B, Firth A. 2015. Insect-Specific Flaviviruses: A Systematic Review of Their Discovery, Host Range, Mode of Transmission, Superinfection Exclusion Potential and Genomic Organization. Viruses 7:1927–1959.

5. Moureau G, Cook S, Lemey P, Nougairède A, Forrester NL, Khasnatinov M, Charrel RN, Firth AE, Gould EA, de Lamballerie X. 2015. New Insights into Flavivirus Evolution, Taxonomy and Biogeographic History, Extended by Analysis of Canonical and Alternative Coding Sequences. PLoS One 10:e0117849--30.

6. Hall RA, Bielefeldt-Ohmann H, McLean BJ, O’Brien CA, Colmant AMG, Piyasena TBH, Harrison JJ, Newton ND, Barnard RT, Prow NA, Deerain JM, Mah MGKY, Hobson-Peters J. 2016. Commensal Viruses of Mosquitoes: Host Restriction, Transmission, and Interaction with Arboviral Pathogens. Evolutionary Bioinformatics 12s2:EBO.S40740.

7. Pierson TC, Diamond MS. 2020. The continued threat of emerging flaviviruses. Nat Microbiol 5:796–812.

8. Porier DL, Wilson SN, Auguste DI, Leber A, Coutermarsh-Ott S, Allen IC, Caswell CC, Budnick JA, Bassaganya-Riera J, Hontecillas R, Weger-Lucarelli J, Weaver SC, Auguste AJ. 2021. Enemy of My Enemy: A Novel Insect-Specific Flavivirus Offers a Promising Platform for a Zika Virus Vaccine. Vaccines (Basel) 9:1142.

9. Hobson-Peters J, Harrison JJ, Watterson D, Hazlewood JE, Vet LJ, Newton ND, Warrilow D, Colmant AMG, Taylor C, Huang B, Piyasena TBH, Chow WK, Setoh YX, Tang B, Nakayama E, Yan K, Amarilla AA, Wheatley S, Moore PR, Finger M, Kurucz N, Modhiran N, Young PR, Khromykh AA, Bielefeldt-Ohmann H, Suhrbier A, Hall RA. 2019. A recombinant platform for flavivirus vaccines and diagnostics using chimeras of a new insect-specific virus. Sci Transl Med 11.

10. Vet LJ, Setoh YX, Amarilla AA, Habarugira G, Suen WW, Newton ND, Harrison JJ, Hobson-Peters J, Hall RA, Bielefeldt-Ohmann H. 2020. Protective Efficacy of a Chimeric Insect-Specific Flavivirus Vaccine against West Nile Virus. Vaccines (Basel) 8:258.

11. Varjak M, Donald CL, Mottram TJ, Sreenu VB, Merits A, Maringer K, Schnettler E, Kohl A. 2017. Characterization of the Zika virus induced small RNA response in Aedes aegypti cells. PLoS Negl Trop Dis 11:e0006010.

12. De Falco L, Silva NM, Santos NC, Huber RG, Martins IC. 2021. The Pseudo-Circular Genomes of Flaviviruses: Structures, Mechanisms, and Functions of Circularization. Cells 10:642.

13. Mazeaud C, Freppel W, Chatel-Chaix L. 2018. The Multiples Fates of the Flavivirus RNA Genome During Pathogenesis. Front Genet 9.

14. Fajardo T, Sanford TJ, Mears H V., Jasper A, Storrie S, Mansur DS, Sweeney TR. 2020. The flavivirus polymerase NS5 regulates translation of viral genomic RNA. Nucleic Acids Res 48:5081–5093.

15. Trettin KD, Sinha NK, Eckert DM, Apple SE, Bass BL. 2017. Loquacious-PD facilitates Drosophila Dicer-2 cleavage through interactions with the helicase domain and dsRNA. Proceedings of the National Academy of Sciences 114:E7939–E7948.

16. Pijlman GP, Funk A, Kondratieva N, Leung J, Torres S, van der Aa L, Liu WJ, Palmenberg AC, Shi PY, Hall RA, Khromykh AA. 2008. A Highly Structured, Nuclease-Resistant, Noncoding RNA Produced by Flaviviruses Is Required for Pathogenicity. Cell Host Microbe 4:579–591.

17. MacFadden A, O’Donoghue Z, Silva PAGCGC, Chapman EG, Olsthoorn RC, Sterken MG, Pijlman GP, Bredenbeek PJ, Kieft JS. 2018. Mechanism and structural diversity of exoribonuclease-resistant RNA structures in flaviviral RNAs. Nat Commun 9:119.

18. Slonchak A, Khromykh AA. 2018. Subgenomic flaviviral RNAs: What do we know after the first decade of research. Antiviral Res 10.1016/j.antiviral.2018.09.006.

19. Vicens Q, Kieft JS. 2021. Shared properties and singularities of exoribonuclease-resistant RNAs in viruses. Comput Struct Biotechnol J 19:4373–4380.

20. Kieft JS, Rabe JL, Chapman EG. 2015. New hypotheses derived from the structure of a flaviviral Xrn1-resistant RNA: Conservation, folding, and host adaptation. RNA Biol 12:1169–1177.

21. Ochsenreiter R, Hofacker I, Wolfinger M. 2019. Functional RNA Structures in the 3′ UTR of Tick-Borne, Insect-Specific and No-Known-Vector Flaviviruses. Viruses 11:298.

22. Villordo SM, Filomatori C V, Sánchez-Vargas I, Blair CD, Gamarnik A V. 2015. Dengue Virus RNA Structure Specialization Facilitates Host Adaptation. PLoS Pathog 11:e1004604--22.

23. Filomatori C V., Carballeda JM, Villordo SM, Aguirre S, Pallarés HM, Maestre AM, Sánchez-Vargas I, Blair CD, Fabri C, Morales MA, Fernandez-Sesma A, Gamarnik A V. 2017. Dengue virus genomic variation associated with mosquito adaptation defines the pattern of viral non-coding RNAs and fitness in human cells. PLoS Pathog 13:e1006265.

24. Funk A, Truong K, Nagasaki T, Torres S, Floden N, Balmori Melian E, Edmonds J, Dong H, Shi PY, Khromykh AA. 2010. RNA structures required for production of subgenomic flavivirus RNA. J Virol 84:11407–11417.

25. Pallarés HM, Costa Navarro GS, Villordo SM, Merwaiss F, de Borba L, Gonzalez Lopez Ledesma MM, Ojeda DS, Henrion-Lacritick A, Morales MA, Fabri C, Saleh MC, Gamarnik A V. 2020. Zika Virus Subgenomic Flavivirus RNA Generation Requires Cooperativity between Duplicated RNA Structures That Are Essential for Productive Infection in Human Cells. J Virol 94.

26. Yeh S-C, Tan W-L, Chowdhury A, Chuo V, Kini RM, Pompon J, Garcia-Blanco MA. 2021. The anti-immune dengue subgenomic flaviviral RNA is found in vesicles in mosquito saliva 2 and associated with increased infectivity. bioRxiv 10.1101/2021.03.02.433543.

27. Moon SL, Anderson JR, Kumagai Y, Wilusz CJ, Akira S, Khromykh AA, Wilusz J. 2012. A noncoding RNA produced by arthropod-borne flaviviruses inhibits the cellular exoribonuclease XRN1 and alters host mRNA stability. RNA 18:2029–2040.

28. Slonchak A, Hugo LE, Freney ME, Hall-Mendelin S, Amarilla AA, Torres FJ, Setoh YX, Peng NYG, Sng JDJ, Hall RA, van den Hurk AF, Devine GJ, Khromykh AA. 2020. Zika virus noncoding RNA suppresses apoptosis and is required for virus transmission by mosquitoes. Nat Commun 11:2205.

29. Hussain M, Torres S, Schnettler E, Funk A, Grundhoff A, Pijlman GP, Khromykh AA, Asgari S. 2012. West Nile virus encodes a microRNA-like small RNA in the 3’ untranslated region which up-regulates GATA4 mRNA and facilitates virus replication in mosquito cells. Nucleic Acids Res 40:2210–2223.

30. Pompon J, Manuel M, Ng GK, Wong B, Shan C, Manokaran G, Soto-Acosta R, Bradrick SS, Ooi EE, Missé D, Shi PY, Garcia-Blanco MA. 2017. Dengue subgenomic flaviviral RNA disrupts immunity in mosquito salivary glands to increase virus transmission. PLoS Pathog 13:e1006535--27.

31. Göertz GP, Fros JJ, Miesen P, Vogels CBF, van der Bent ML, Geertsema C, Koenraadt CJM, van Rij RP, van Oers MM, Pijlman GP. 2016. Noncoding Subgenomic Flavivirus RNA Is Processed by the Mosquito RNA Interference Machinery and Determines West Nile Virus Transmission by Culex pipiens Mosquitoes. J Virol 90:10145–10159.

32. Moon SL, Dodd BJT, Brackney DE, Wilusz CJ, Ebel GD, Wilusz J. 2015. Flavivirus sfRNA suppresses antiviral RNA interference in cultured cells and mosquitoes and directly interacts with the RNAi machinery. Virology 485:322–329.

33. Phoo WW, Li Y, Zhang Z, Lee MY, Loh YR, Tan YB, Ng EY, Lescar J, Kang C, Luo D. 2016. Structure of the NS2B-NS3 protease from Zika virus after self-cleavage. Nat Commun 1–8.

34. Schnettler E, Sterken MG, Leung JY, Metz SW, Geertsema C, Goldbach RW, Vlak JM, Kohl A, Khromykh AA, Pijlman GP. 2012. Noncoding Flavivirus RNA Displays RNA Interference Suppressor Activity in Insect and Mammalian Cells. J Virol 86:13486– 13500.

35. Cirimotich CM, Scott JC, Phillips AT, Geiss BJ, Olson KE. 2009. Suppression of RNA interference increases alphavirus replication and virus-associated mortality in Aedes aegypti mosquitoes. BMC Microbiol 9:49.

36. Bronkhorst AW, van Rij RP. 2014. The long and short of antiviral defense: small RNA-based immunity in insects. Curr Opin Virol 7:19–28.

37. Marques JT, Imler J-L. 2016. The diversity of insect antiviral immunity: insights from viruses. Curr Opin Microbiol 32:71–76.

38. Rosendo Machado S, van der Most T, Miesen P. 2021. Genetic determinants of antiviral immunity in dipteran insects – Compiling the experimental evidence. Dev Comp Immunol 119:104010.

39. Samuel GH, Pohlenz T, Dong Y, Coskun N, Adelman ZN, Dimopoulos G, Myles KM. 2023. RNA interference is essential to modulating the pathogenesis of mosquito-borne viruses in the yellow fever mosquito *Aedes aegypti*. Proceedings of the National Academy of Sciences 120.

40. Dong S, Dimopoulos G. 2023. Aedes aegypti Argonaute 2 controls arbovirus infection and host mortality. Nat Commun 14:5773.

41. Merkling SH, Crist AB, Henrion-Lacritick A, Frangeul L, Couderc E, Gausson V, Blanc H, Bergman A, Baidaliuk A, Romoli O, Saleh M-C, Lambrechts L. 2023. Multifaceted contributions of Dicer2 to arbovirus transmission by Aedes aegypti. Cell Rep 42:112977.

42. Elrefaey AME, Hollinghurst P, Reitmayer CM, Alphey L, Maringer K. 2021. Innate Immune Antagonism of Mosquito-Borne Flaviviruses in Humans and Mosquitoes. Viruses 13:2116.

43. Miesen P, Joosten J, van Rij RP. 2016. PIWIs Go Viral: Arbovirus-Derived piRNAs in Vector Mosquitoes. PLoS Pathog 12:e1006017.

44. Blair CD. 2019. Deducing the Role of Virus Genome-Derived PIWI-Associated RNAs in the Mosquito–Arbovirus Arms Race. Front Genet 10.

45. Miesen P, Ivens A, Buck AH, van Rij RP. 2016. Small RNA Profiling in Dengue Virus 2-Infected Aedes Mosquito Cells Reveals Viral piRNAs and Novel Host miRNAs. PLoS Negl Trop Dis 10:e0004452.

46. Miesen P, Girardi E, van Rij RP. 2015. Distinct sets of PIWI proteins produce arbovirus and transposon-derived piRNAs in Aedes aegypti mosquito cells. Nucleic Acids Res 43:6545–6556.

47. Joosten J, Miesen P, Ta\c sköprü E, Pennings B, Jansen PWTC, Huynen MA, Vermeulen M, van Rij RP. 2019. The Tudor protein Veneno assembles the ping-pong amplification complex that produces viral piRNAs in Aedes mosquitoes. Nucleic Acids Res 47:2546–2559.

48. Varjak M, Donald CL, Mottram TJ, Sreenu VB, Merits A, Maringer K, Schnettler E, Kohl A. 2017. Characterization of the Zika virus induced small RNA response in Aedes aegypti cells. PLoS Negl Trop Dis 11:e0006010.

49. Tassetto M, Kunitomi M, Whitfield ZJ, Dolan PT, Sánchez-Vargas I, Garcia-Knight M, Ribiero I, Chen T, Olson KE, Andino R. 2019. Control of RNA viruses in mosquito cells through the acquisition of vDNA and endogenous viral elements. Elife 8.

50. Suzuki Y, Baidaliuk A, Miesen P, Frangeul L, Crist AB, Merkling SH, Fontaine A, Lequime S, Moltini-Conclois I, Blanc H, van Rij RP, Lambrechts L, Saleh M-C. 2020. Non-retroviral Endogenous Viral Element Limits Cognate Virus Replication in Aedes aegypti Ovaries. Current Biology 30:3495–3506.e6.

51. Palatini U, Miesen P, Carballar-Lejarazu R, Ometto L, Rizzo E, Tu Z, van Rij RP, Bonizzoni M. 2017. Comparative genomics shows that viral integrations are abundant and express piRNAs in the arboviral vectors Aedes aegypti and Aedes albopictus. BMC Genomics 18:512.

52. Crava CM, Varghese FS, Pischedda E, Halbach R, Palatini U, Marconcini M, Gasmi L, Redmond S, Afrane Y, Ayala D, Paupy C, Carballar-Lejarazu R, Miesen P, Rij RP, Bonizzoni M. 2021. Population genomics in the arboviral vector Aedes aegypti reveals the genomic architecture and evolution of endogenous viral elements. Mol Ecol 30:1594–1611.

53. Crabtree MB, Sang RC, Stollar V, Dunster LM, Miller BR. 2003. Genetic and phenotypic characterization of the newly described insect flavivirus, Kamiti River virus. Arch Virol 148:1095–1118.

54. Kopp A, Gillespie TR, Hobelsberger D, Estrada A, Harper JM, Miller RA, Eckerle I, Müller MA, Podsiadlowski L, Leendertz FH, Drosten C, Junglen S. 2013. Provenance and Geographic Spread of St. Louis Encephalitis Virus. mBio 4.

55. Göertz G, Miesen P, Overheul G, van Rij R, van Oers M, Pijlman G. 2019. Mosquito Small RNA Responses to West Nile and Insect-Specific Virus Infections in Aedes and Culex Mosquito Cells. Viruses 11:271.

56. Saiyasombat R, Bolling BG, Brault AC, Bartholomay LC, Blitvich BJ. 2011. Evidence of Efficient Transovarial Transmission of Culex Flavivirus by Culex pipiens (Diptera: Culicidae). J Med Entomol 48:1031–1038.

57. Rozen-Gagnon K, Gu M, Luna JM, Luo J-D, Yi S, Novack S, Jacobson E, Wang W, Paul MR, Scheel TKH, Carroll T, Rice CM. 2021. Argonaute-CLIP delineates versatile, functional RNAi networks in Aedes aegypti, a major vector of human viruses. Cell Host Microbe 29:834–848.e13.

58. Wang Y, Jin B, Liu P, Li J, Chen X, Gu J. 2018. piRNA Profiling of Dengue Virus Type 2-Infected Asian Tiger Mosquito and Midgut Tissues. Viruses 10:213.

59. MacFadden A, O’Donoghue Z, Silva PAGC, Chapman EG, Olsthoorn RC, Sterken MG, Pijlman GP, Bredenbeek PJ, Kieft JS. 2018. Mechanism and structural diversity of exoribonuclease-resistant RNA structures in flaviviral RNAs. Nat Commun 9:119.

60. Slonchak A, Parry R, Pullinger B, Sng JDJ, Wang X, Buck TF, Torres FJ, Harrison JJ, Colmant AMG, Hobson-Peters J, Hall RA, Tuplin A, Khromykh AA. 2022. Structural analysis of 3’UTRs in insect flaviviruses reveals novel determinants of sfRNA biogenesis and provides new insights into flavivirus evolution. Nat Commun 13:1279.

61. Göertz GP, van Bree JWM, Hiralal A, Fernhout BM, Steffens C, Boeren S, Visser TM, Vogels CBF, Abbo SR, Fros JJ, Koenraadt CJM, van Oers MM, Pijlman GP. 2019. Subgenomic flavivirus RNA binds the mosquito DEAD/H-box helicase ME31B and determines Zika virus transmission by Aedes aegypti. Proceedings of the National Academy of Sciences 116:19136–19144.

62. Gritsun DJ, Jones IM, Gould EA, Gritsun TS. 2014. Molecular archaeology of Flaviviridae untranslated regions: duplicated RNA structures in the replication enhancer of flaviviruses and pestiviruses emerged via convergent evolution. PLoS One 9:e92056.

63. Łabno A, Tomecki R, Dziembowski A. 2016. Cytoplasmic RNA decay pathways - Enzymes and mechanisms. Biochimica et Biophysica Acta (BBA) - Molecular Cell Research 1863:3125–3147.

64. Chen Y-S, Fan Y-H, Tien C-F, Yueh A, Chang R-Y. 2018. The conserved stem-loop II structure at the 3’ untranslated region of Japanese encephalitis virus genome is required for the formation of subgenomic flaviviral RNA. PLoS One 13:e0201250--17.

65. Akiyama BM, Laurence HM, Massey AR, Costantino DA, Xie X, Yang Y, Shi PY, Nix JC, Beckham JD, Kieft JS. 2016. Zika virus produces noncoding RNAs using a multi-pseudoknot structure that confounds a cellular exonuclease. Science (1979) 10.1126/science.aah3963.

66. Bolling B, Weaver S, Tesh R, Vasilakis N. 2015. Insect-Specific Virus Discovery: Significance for the Arbovirus Community. Viruses 7:4911–4928.

67. Qiu Y, Xu Y-P, Wang M, Miao M, Zhou H, Xu J, Kong J, Zheng D, Li R-T, Zhang R-R, Guo Y, Li X-F, Cui J, Qin C-F, Zhou X. 2020. Flavivirus induces and antagonizes antiviral RNA interference in both mammals and mosquitoes. Sci Adv 6.

68. Blair C, Olson K. 2015. The Role of RNA Interference (RNAi) in Arbovirus-Vector Interactions. Viruses 7:820–843.

69. Lee W-S, Webster JA, Madzokere ET, Stephenson EB, Herrero LJ. 2019. Mosquito antiviral defense mechanisms: a delicate balance between innate immunity and persistent viral infection. Parasit Vectors 12:165.

70. Colmant AMG, Hobson-Peters J, Bielefeldt-Ohmann H, van den Hurk AF, Hall-Mendelin S, Chow WK, Johansen CA, Fros J, Simmonds P, Watterson D, Cazier C, Etebari K, Asgari S, Schulz BL, Beebe N, Vet LJ, Piyasena TBH, Nguyen H-D, Barnard RT, Hall RA. 2017. A New Clade of Insect-Specific Flaviviruses from Australian Anopheles Mosquitoes Displays Species-Specific Host Restriction. mSphere 2.

71. Rückert C, Prasad AN, Garcia-Luna SM, Robison A, Grubaugh ND, Weger-Lucarelli J, Ebel GD. 2019. Small RNA responses of Culex mosquitoes and cell lines during acute and persistent virus infection. Insect Biochem Mol Biol 109:13–23.

72. Fan Y-H, Nadar M, Chen C-C, Weng C-C, Lin Y-T, Chang R-Y. 2011. Small Noncoding RNA Modulates Japanese Encephalitis Virus Replication and Translation in trans. Virol J 8:492.

73. Alvarez DE, Lodeiro MF, Ludueña SJ, Pietrasanta LI, Gamarnik A V. 2005. Long-Range RNA-RNA Interactions Circularize the Dengue Virus Genome. J Virol 79:6631– 6643.

74. Roby J, Pijlman G, Wilusz J, Khromykh A. 2014. Noncoding Subgenomic Flavivirus RNA: Multiple Functions in West Nile Virus Pathogenesis and Modulation of Host Responses. Viruses 6:404–427.

75. Clarke DK, Duarte EA, Elena SF, Moya A, Domingo E, Holland J. 1994. The red queen reigns in the kingdom of RNA viruses. Proceedings of the National Academy of Sciences 91:4821–4824.

76. Junglen S, Kopp A, Kurth A, Pauli G, Ellerbrok H, Leendertz FH. 2009. A New Flavivirus and a New Vector: Characterization of a Novel Flavivirus Isolated from Uranotaenia Mosquitoes from a Tropical Rain Forest. J Virol 83:4462–4468.

77. Bassett AR, Tibbit C, Ponting CP, Liu J-L. 2014. Mutagenesis and homologous recombination in Drosophila cell lines using CRISPR/Cas9. Biol Open 3:42–49.

78. Afgan E, Baker D, Batut B, van den Beek M, Bouvier D, Čech M, Chilton J, Clements D, Coraor N, Grüning BA, Guerler A, Hillman-Jackson J, Hiltemann S, Jalili V, Rasche H, Soranzo N, Goecks J, Taylor J, Nekrutenko A, Blankenberg D. 2018. The Galaxy platform for accessible, reproducible and collaborative biomedical analyses: 2018 update. Nucleic Acids Res 46:W537–W544.

79. de Borba L, Villordo SM, Marsico FL, Carballeda JM, Filomatori C V., Gebhard LG, Pallarés HM, Lequime S, Lambrechts L, Sánchez Vargas I, Blair CD, Gamarnik A V. 2019. RNA Structure Duplication in the Dengue Virus 3′ UTR: Redundancy or Host Specificity? mBio 10.

80. Lorenz R, Bernhart SH, Höner zu Siederdissen C, Tafer H, Flamm C, Stadler PF, Hofacker IL. 2011. ViennaRNA Package 2.0. Algorithms for Molecular Biology 6:26.

81. Katoh K, Standley DM. 2013. MAFFT Multiple Sequence Alignment Software Version 7: Improvements in Performance and Usability. Mol Biol Evol 30:772–780.

82. Criscuolo A, Gribaldo S. 2010. BMGE (Block Mapping and Gathering with Entropy): a new software for selection of phylogenetic informative regions from multiple sequence alignments. BMC Evol Biol 10:210.

83. Guindon S, Dufayard J-F, Lefort V, Anisimova M, Hordijk W, Gascuel O. 2010. New Algorithms and Methods to Estimate Maximum-Likelihood Phylogenies: Assessing the Performance of PhyML 3.0. Syst Biol 59:307–321.

84. Lemoine F, Correia D, Lefort V, Doppelt-Azeroual O, Mareuil F, Cohen-Boulakia S, Gascuel O. 2019. NGPhylogeny.fr: new generation phylogenetic services for non-specialists. Nucleic Acids Res 47:W260–W265.

85. Letunic I, Bork P. 2021. Interactive Tree Of Life (iTOL) v5: an online tool for phylogenetic tree display and annotation. Nucleic Acids Res 49:W293–W296.

86. Brown NP, Leroy C, Sander C. 1998. MView: a web-compatible database search or multiple alignment viewer. Bioinformatics 14:380–381.

87. Nicholson J, Ritchie SA, Van Den Hurk AF. 2014. Aedes albopictus (Diptera: Culicidae) as a Potential Vector of Endemic and Exotic Arboviruses in Australia. J Med Entomol 51:661–669.

