## Supplemental files for "Pan-flavivirus analysis reveals sfRNA-independent, 3’UTR-biased siRNA production from an Insect-Specific Flavivirus"

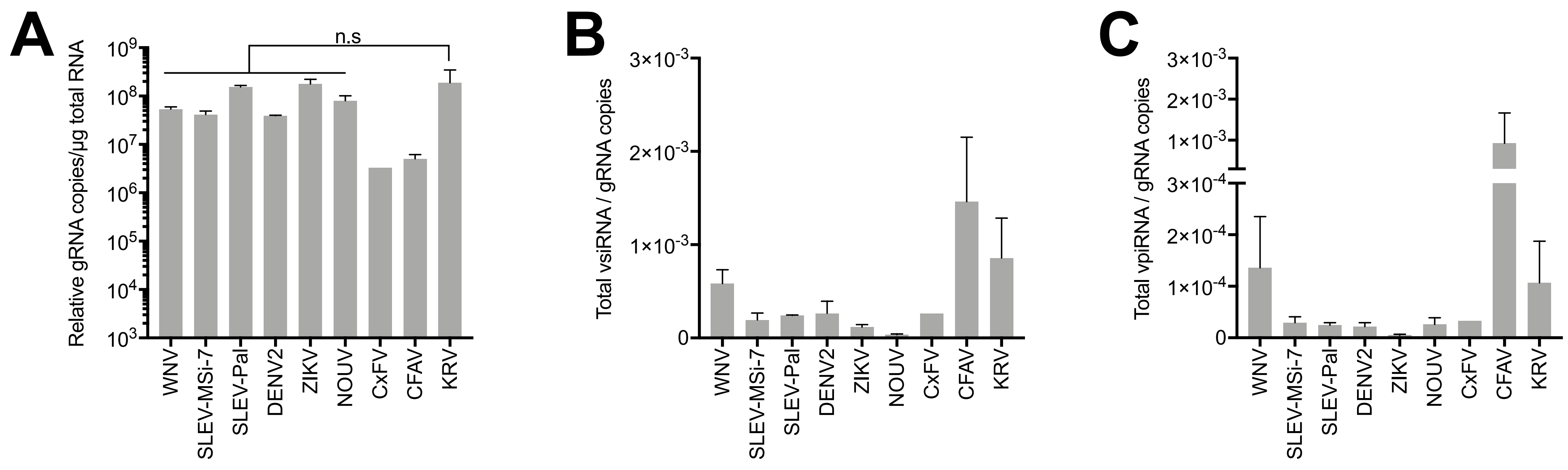
A
Relative gRNA copies/µg total RNA
n.s
WNV
SLEV-MSi-7
SLEV-Pal
DENV2
ZIKV
NOUV
CxFV
CFAV
KRV
B
Total vsiRNA / gRNA copies
C
Total vpiRNA / gRNA copies

### Fig. S1

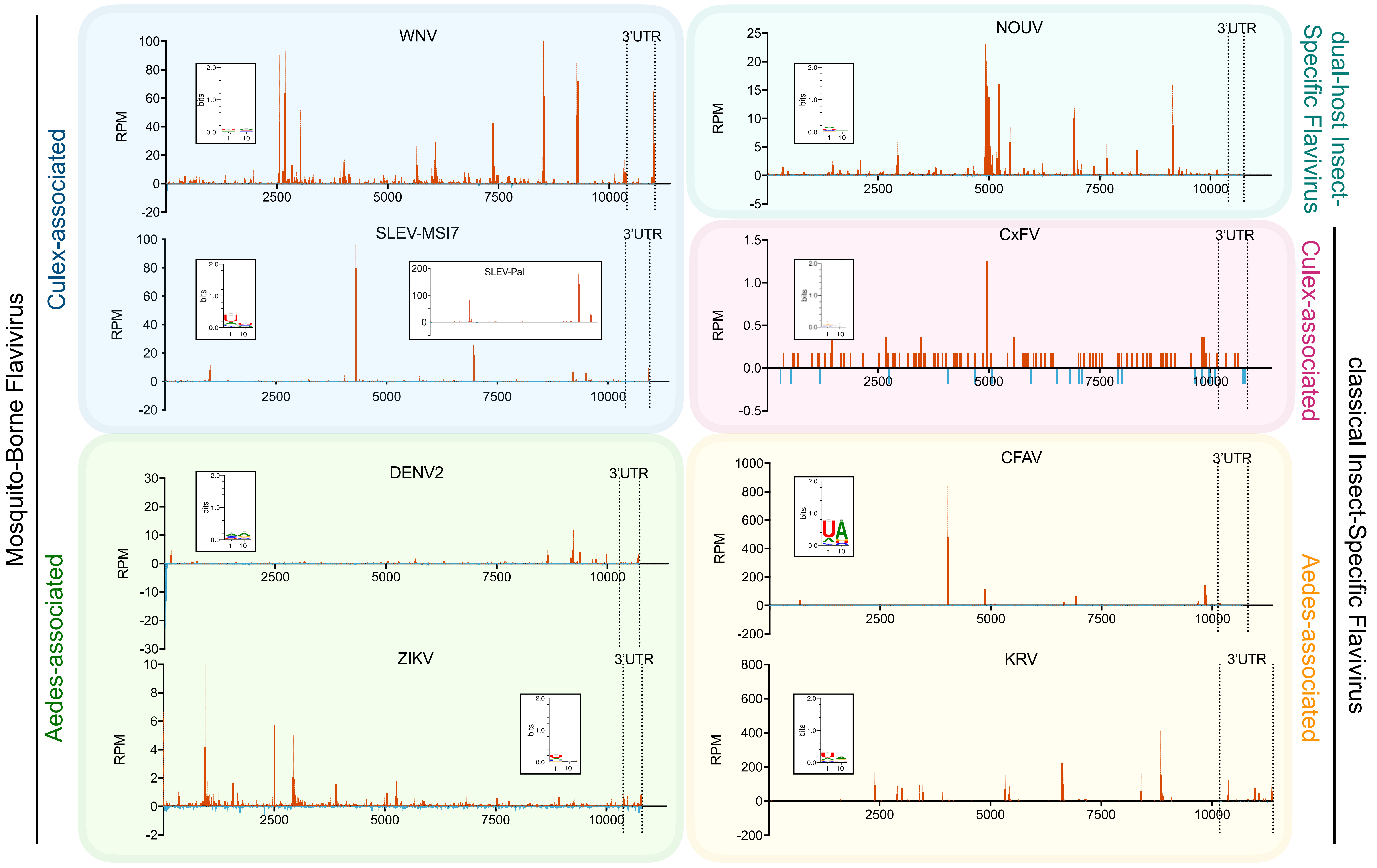

Mosquito-Borne Flavivirus
Culex-associated
Aedes-associated
WNV
SLEV-MSI7
SLEV-Pal
DENV2
ZIKV
NOUV
CxFV
CFAV
KRV
3'UTR
RPM
bits
2500
5000
7500
10000
dual-host Insect-Specific Flavivirus
classical Insect-Specific Flavivirus
Culex-associated
Aedes-associated


Fig. S2

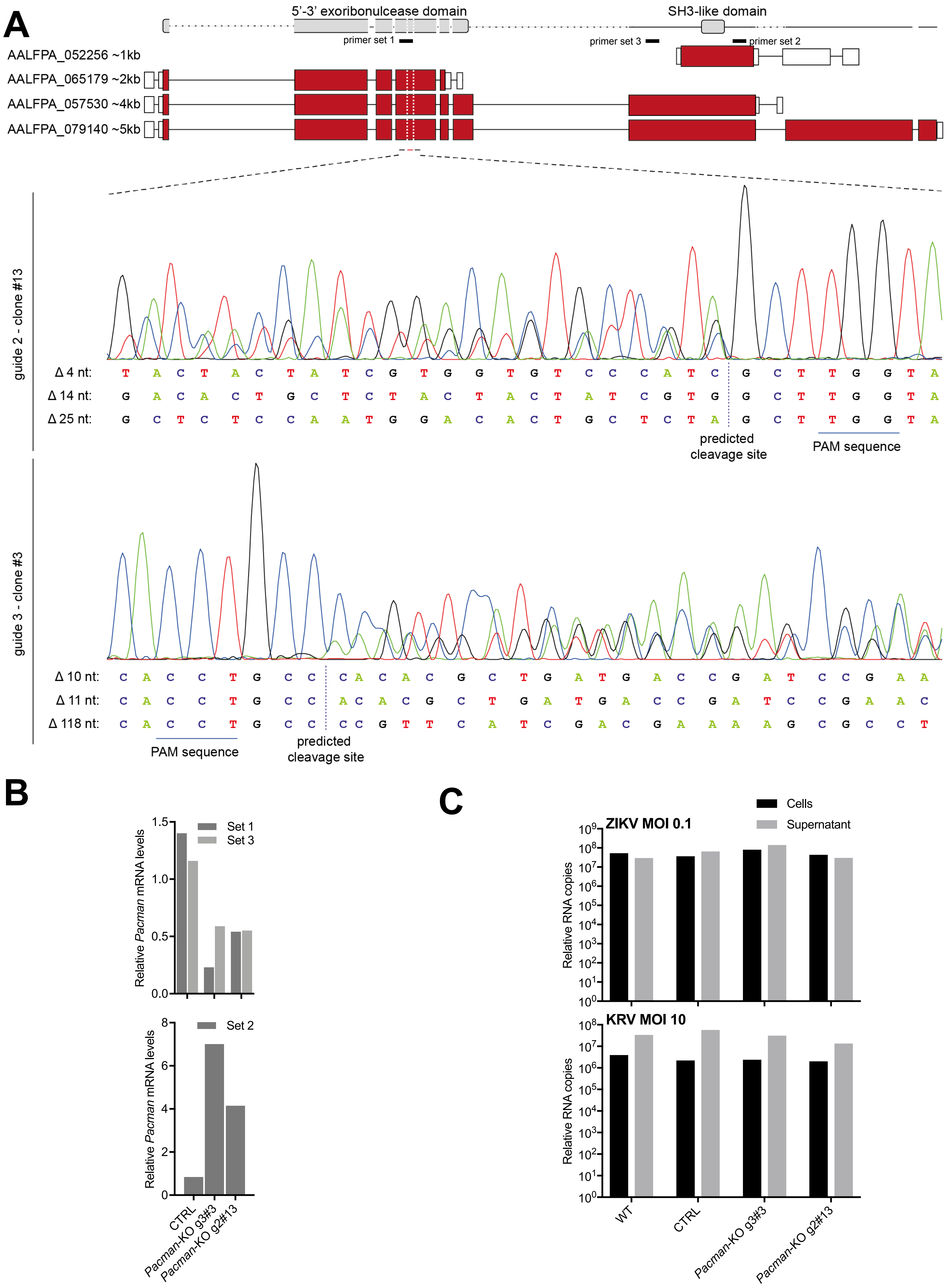
A
5'-3' exoribonulcease domain
SH3-like domain
primer set 1
primer set 3
primer set 2
AALFPA_052256 ~1kb
AALFPA_065179 ~2kb
AALFPA_057530 ~4kb
AALFPA_079140 ~5kb
guide 2 - clone #13
Δ 4 nt: T A C T A C T A T C G T G G T G T C C C A T C G C T T G G T A
Δ 14 nt: G A C A C T G C T C T A C T A C T A T C G T G G C T T G G T A
Δ 25 nt: G C T C T C C A A T G G A C A C T G C T C T A G C T T G G T A
predicted cleavage site
PAM sequence
guide 3 - clone #3
Δ 10 nt: C A C C T G C C C A C A C G C T G A T G A C C G A T C C G A A
Δ 11 nt: C A C C T G C C A C A C G C T G A T G A C C G A T C C G A A C
Δ 118 nt: C A C C T G C C C C G T T C A T C G A C G A A A A G C G C C T
PAM sequence
predicted cleavage site
B
Set 1
Set 3
Relative Pacman mRNA levels
Set 2
CTRL
Pacman-KO g3#3
Pacman-KO g2#13
C
Cells
Supernatant
ZIKV MOI 0.1
KRV MOI 10
Relative RNA copies
WT
CTRL
Pacman-KO g3#3
Pacman-KO g2#13

### Fig. S3

**Supplementary table 1. List of viruses used in pan-flavivirus small RNA analysis**

| Virus | Strain/Isolate | Start CDS | End CDS | 3'UTR start | End | Reference |
|---|---|---|---|---|---|---|
| CFAV | RioPiedras | 104 | 10129 | 10127 | 10682 | NC_001564.2 |
| CxFV | Uganda08 | 92 | 10183 | 10181 | 10837 | NC_008604.2 |
| DENV2 | 16881 | 97 | 10272 | 10270 | 10723 | NC_001474.2 |
| KRV | SR-75 | 97 | 10170 | 10168 | 11375 | NC_005064.1 |
| NOUV | B3 | 80 | 10408 | 10406 | 10755 | NC_033715.1 |
| SLEV | Palenque | 99 | 10391 | 10389 | 10938 | JQ957869.1 |
| SLEV | MSI-7 | 99 | 10391 | 10389 | 10939 | MSI-7(incomplete): DQ359217.1; Hubbard (closest complete): EU566860.1, Refseq: NC_007580.2 |
| WNV | NY99 | 97 | 10398 | 10396 | 11029 | NC_009942.1 |
| ZIKV | H/PF/2013 | 108 | 10379 | 10377 | 10807 | KJ776791.2 |

**Supplementary table 2. List of oligonucleotides for northern blots, qPCR and cloning**

| Use | Virus | Probe sequence | Position | Notes |
|---|---|---|---|---|
| Northern Blot Probes | KRV (+)RNA | CAGGGGACTCGGGGGAGCGGGT | 10392, 10988 | - |
| | | GAGCGGGTGCGTCASGCCCGACAC | 10972 | - |
| | | TATCTTTTCTATACCATARATGC | 11350 | - |
| | KRV (-)RNA | ACCCGCTCCCCCGAGTCCCCTG | 10392, 10988 | - |
| | | CTGGTTCTCGCAACTCCAGTCGAA | 10599 | - |
| | | GTGTCGGGCSTGACGCACCCGCTC | 10972 | - |
| | | TATCTTTTCTATACCATARATGC | 11350 | - |
| qPCR primers | KRV gRNA For | GGTCAATGAGACCGAACGA | 7602 | - |
| | KRV gRNA Rev | GTGTATCCATACACAGACGAC | 7758 | - |
| | KRV sfRNA1 For | GGTGACCTGTCTCATACATG | 10503 | - |
| | KRV sfRNA1 Rev | ACATTGCTGATCCTTGTTCC | 10637 | - |
| | KRV sfRNA2 For | GTCATAGGCACCTGACCTG | 11073 | - |
| | KRV sfRNA2 Rev | TCCGTCCGGTTTTGAAAGC | 11232 | - |
| | Aalb pacman KO For | GCAACTCGGGCCGAACAAAC | 1321 | Sanger sequencing of Cas9 edited region |
| | Aalb pacman KO Rev | CTGATCGCGATAAATTGCCG | 1975 | |
| | Aalb pacman Set1 For | GGGCTACCAGGACTATAATG | 1299 | AALFPA-065179 / 057530 / 079140 |
| | Aalb pacman Set1 Rev | TCCCGTTGAGATCGGTTTCG | 1639 | |
| | Aalb pacman Set2 For | CAGAGGAAGCACGAGTTGGT | 3568 | AALFPA-052256 / 057530 / 079140 |
| | Aalb pacman Set2 Rev | ATGTGGACGGCACTTGTGAT | 3802 | |
| | Aalb pacman Set3 For | GCGCTGAAAACTCTGAAGCC | 2431 | AALFPA-057530 / 079140 |
| | Aalb pacman Set3 Rev | CCAGATTGGGTTCCTCGTGT | 2602 | |
| | RPL5 For | TCGCTTACGCCCGCATTGAGGGTGAT | - | Housekeeping gene |
| | RPL5 Rev | TCGCCGGTCACATCGGTACAGCCA | - | |
| Cloning of viral sequences | WNV NS5 For | TGAAGAGCCCCAACTAGTGC | 8010 | - |
| | WNV NS5 Rev | TTCAAGGACCCGAATCGTCC | 8159 | |

| | | | | |
|---|---|---|---|---|
| | SLEV-MSI7 NS5 For | GAGAGAAGGGCGTCTCACAG | 7802 | - |
| | SLEV-MSI7 NS5 Rev | GAACATGCTTCAGGGTTGCG | 7943 | |
| | SLEV-Pal NS5 For | CACGTCCAAGAGGTGAAGGG | 7956 | - |
| | SLEV-Pal NS5 Rev | TCACAGCTCGGGTTTGACTC | 8115 | |
| | DENV2 NS5 For | GTAGTGGACCTCGGTTGTGG | 7798 | - |
| | DENV2 NS5 Rev | GTGTCACACTTTTCTGGCGG | 7975 | |
| | ZIKV NS3 For | GAGAGAGTCATTCTGGCTGGA | 5925 | - |
| | ZIKV NS3 Rev | TCCCTCAATGGCTGCTACTT | 6154 | |
| | NOUV NS5 For | AAGCCTACACGAAAGGAGGC | 7992 | - |
| | NOUV NS5 Rev | ACGACTCCCCAATGTCACAC | 8122 | |
| | CxFV NS1 For | GATCCGGAGGGTTTGTGTGG | 2265 | - |
| | CxFV NS1 Rev | GCATTGTAGGACATCCTCAC | 2400 | |
| | CFAV NS1 For | GCAGCGGCGCTTTTGTGTGG | 2265 | - |
| | CFAV NS1 Rev | GCACTGCAAGGCATCCTCAC | 2400 | |
| | KRV NS5 For | ATCCACAGCTGTAGGCCTTG | 7515 | - |
| | KRV NS5 Rev | CAACCCGTCCGTTTGGTTTC | 7680 | |
| | sfRNA1 For | CGACTCTAGAGGATCC-GTATGAGACAGGTCACCACT | 1126 | - |
| | sfRNA1 Rev | ATTCGAGCTCGGTACC-GCAATGCACTCCTGAGTAG | 10500 | |
| | sfRNA2 For | CGACTCTAGAGGATCC-TAAGGCGCCACTCTTATCC | 1126 | - |
| | sfRNA2 Rev | ATTCGAGCTCGGTACC-GCAATGCACTCCTGAGTAG | 11092 | |
| | sfRNA1' and 2' For | CGACTCTAGAGGATCC-ATGGCGTTTTCAATGAGATAGG | 1126 | - |
| | sfRNA1' and 2' Rev | ATTCGAGCTCGGTACC-GCAATGCACTCCTGAGTAG | 10933 | |
| **Cloning of guide RNA for CRISPR/Cas9 gene editing** | Amp till tracrRNA (XbaI) For | TCCTTCGGTCCTCCGATCGTTG | - | In-Fusion #1 |
| | Amp till tracrRNA (XbaI) Rev | TGCTTTTTTTCTAGAAGATCTGGAAAAATGATGTG | | |
| | tracrRNA till Aalb pU6 (XbaI) For | TCTAGAAAAAAAGCACCGACTCGGTG | - | In-Fusion #2 |
| | tracrRNA till Aalb pU6 (XbaI) Rev | GACGGCAATGAAATGGAAGAGCGAGCTCTTCC | | |
| | Aalb pU6-2 till pAc (XbaI) For | CATTTCATTGCCGTCGCTTC | - | In-Fusion #3 |

| | | | | |
|---|---|---|---|---|
| | Aalb pU6-2 till pAc (XbaI) Rev | CGAGATCTGTCTAGACTCAGCTCGAGTGTGGTCTTAG | | |
| | pAc Fw (XbaI) For | TCTAGACAGATCTCGCTGCCTGTTATG | - | In-Fusion #4 |
| | pAc Fw (XbaI) Rev | GCCAAGAATGGAGCGATCGC | | |
| | Aalb pacman guide #2.1 For | AATGTGGTGTCCCATCCTGGGCT | - | - |
| | Aalb pacman guide #2.1 Rev | AACAGCCCAGGATGGGACACCAC | | |
| | Aalb pacman guide #3.1 For | AATGCGTGTGGTAAGCACTGGGC | - | - |
| | Aalb pacman guide #3.1 Rev | AACGCCCAGTGCTTACCACACGC | | |

**Supplementary table 3. List of genome references used for 3'UTR analysis**

| RefSeq | Clade | Full name | Abbreviation | 5'UTR size (nt) | CDS size (nt) | 3'UTR size (nt) |
|---|---|---|---|---|---|---|
| NC_012932.1 | ISFV | Aedes flavivirus | AEFV | 96 | 10026 | 942 |
| NC_001564.2 | ISFV | Cell fusing agent virus | CFAV | 103 | 10026 | 553 |
| NC_008604.2 | ISFV | Culex flavivirus | CxFV | 91 | 10092 | 654 |
| NC_005064.1 | ISFV | Kamiti River virus | KRV | 96 | 10074 | 1208 |
| NC_027819.1 | ISFV | Mercadeo virus | MECDV | 88 | 10212 | 638 |
| NC_021069.1 | ISFV | Mosquito flavivirus | MSFV | 111 | 10080 | 674 |
| NC_027817.1 | ISFV | Parramatta River virus | PaRV | 109 | 10155 | 629 |
| NC_012671.1 | ISFV | Quang Binh virus | QBV | 112 | 10080 | 673 |
| NC_017086.1 | ISFV | Chaoyang virus | CHAOV | 99 | 10308 | 326 |
| NC_016997.1 | ISFV | Donggang virus | DONV | 113 | 10335 | 343 |
| NC_027999.1 | ISFV | Paraiso Escondido virus | EPEV | 119 | 10326 | 316 |
| NC_040610.1 | ISFV | Nanay virus | NANV | 106 | 10299 | 399 |
| NC_034017.1 | ISFV | Xishuangbanna aedes flavivirus | XFV | 90 | 10245 | 549 |
| NC_009026.2 | MBFV | Aroa virus | AROAV | 104 | 10290 | 421 |
| NC_001477.1 | MBFV | Dengue virus 1 | DENV1 | 94 | 10179 | 462 |
| NC_001474.2 | MBFV | Dengue virus 2 | DENV2 | 96 | 10176 | 451 |
| NC_001475.2 | MBFV | Dengue virus 3 | DENV3 | 94 | 10173 | 440 |
| NC_002640.1 | MBFV | Dengue virus 4 | DENV4 | 101 | 10164 | 384 |
| NC_012533.1 | MBFV | Kedougou virus | KEDV | 106 | 10227 | 390 |
| NC_001437.1 | MBFV | Japanese encephalitis virus | JEV | 95 | 10299 | 582 |
| NC_000943.1 | MBFV | Murray Valley encephalitis virus | MVEV | 95 | 10305 | 614 |
| NC_007580.2 | MBFV | Saint Louis encephalitis virus | SLEV | 98 | 10293 | 549 |
| NC_009942.1 | MBFV | West Nile virus | WNV lin.1 | 96 | 10302 | 631 |
| NC_001563.2 | MBFV | West Nile virus | WNV lin.2 | 96 | 10293 | 573 |
| NC_009029.2 | MBFV | Kokobera virus | KOKV | 83 | 10233 | 558 |
| NC_012534.1 | MBFV | Bagaza virus | BAGV | 94 | 10281 | 566 |
| NC_009028.2 | MBFV | Ilheus virus | ILHV | 92 | 10275 | 388 |
| NC_040776.1 | MBFV | Rocio virus | ROCV | 92 | 10278 | 424 |
| NC_034151.1 | MBFV | T'Ho virus | THOV | 97 | 10284 | 556 |
| NC_035889.1 | MBFV | Zika virus | ZIKV/2015 | 107 | 10272 | 429 |
| NC_012532.1 | MBFV | Zika virus | ZIKV/1947 | 106 | 10260 | 428 |
| NC_008719.1 | MBFV | Sepik virus | SEPV | 116 | 10218 | 459 |
| NC_012735.1 | MBFV | Wesselsbron virus | WESSV | 118 | 10218 | 478 |
| NC_002031.1 | MBFV | Yellow fever virus 17D | YFV17D | 118 | 10236 | 508 |
| NC_005039.1 | NKV | Yokose virus | YOKV | 150 | 10278 | 429 |
| NC_003635.1 | NKV | Modoc virus | MODV | 109 | 10125 | 366 |
| NC_004119.1 | NKV | Montana myotis leukoencephalitis virus | MMLV | 108 | 10125 | 460 |
| NC_003690.1 | TBFV | Langat virus | LGTV | 130 | 10245 | 568 |

| | | | | | | |
|---|---|---|---|---|---|---|
| NC_001809.1 | TBFV | Louping ill virus | LIV | 129 | 10245 | 500 |
| NC_005062.1 | TBFV | Omsk hemorrhagic fever virus | OHFV | 132 | 10245 | 410 |
| NC_003687.1 | TBFV | Powassan virus | POWV | 111 | 10248 | 480 |
| NC_027709.1 | TBFV | Spanish goat encephalitis virus | SGEV | 132 | 10245 | 493 |
| NC_001672.1 | TBFV | Tick-borne encephalitis virus | TBEV | 132 | 10245 | 764 |
| | | | | | **Median** | 493 |
